# THE COSTS AND BENEFITS OF BASAL INFECTION RESISTANCE VS DIVERSE IMMUNE PRIMING RESPONSES IN AN INSECT

**DOI:** 10.1101/734038

**Authors:** Arun Prakash, Deepa Agashe, Imroze Khan

## Abstract

In insects, basal pathogen resistance and immune priming can evolve as mutually exclusive strategies, with distinct infection outcomes. However, the evolutionary drivers of such diverse immune functions remain poorly understood. Here, we addressed this key issue by systematically analyzing the differential fitness costs and benefits of priming vs. resistance evolution in *Tribolium* beetle populations infected with *Bacillus thuringiensis*. Surprisingly, resistant beetles had increased post-infection reproduction and a longer lifespan under both starving as well as fed conditions, with no other measurable costs. In contrast, priming reduced offspring early survival, development rate and reproduction. Priming did improve post-infection survival of offspring, but this added trans-generational benefit of immune priming might not compensate for its pervasive costs. Resistance was thus consistently more beneficial. Our work represents the first report of experimentally evolved trans-generational priming, and a detailed comparison of the complex fitness consequences of evolved priming vs resistance.

**HIGHLIGHTS:**

1. Divergent costs and benefits of experimentally evolved immune priming vs resistance
2. Increased reproduction and lifespan in resistant populations
3. No other hidden costs of resistance
4. In contrast, reduced juvenile fitness and reproduction in primed populations
5. First evidence for experimentally evolved trans-generational immune priming

## 1. INTRODUCTION

Until recently, it was assumed that insects have nonspecific immunity and cannot build immune memory against previously encountered pathogens, since they lack the immune cells responsible for adaptive immunity in vertebrates (Cooper and Eleftherianos, 2017). Now, growing evidence contradicts this traditional view: in addition to basal resistance mechanisms against infections (Rolff and Siva-Jothy, 2003), priming with a sub-lethal exposure to a pathogen often protects against subsequent exposure to the same pathogen. This survival benefit of priming is observed both in later life stages of primed individuals (“within-generation immune priming”; henceforth WGIP), and in their offspring (“trans-generational immune priming”; henceforth TGIP), in a range of insect species (reviewed in Milutinović et al., 2016) including Dipterans (Pham et al., 2007; Ramirez et al., 2015; Ramirez et al., 2017), Coleopterans (Roth et al., 2009; Roth et al., 2010; Khan et al., 2016), Lepidopterans (Fallon et al., 2011) and Hymenopterans (Sadd and Schmid-Hempel, 2006). Theoretical studies also highlight the importance of priming in reducing infection prevalence and regulating population size, stability and age structure during infection (Tate and Rudolf, 2012; Best et al., 2013). Thus, it appears that under pathogen pressure, priming responses should be selectively favoured. Recently, we also, for the first time, directly demonstrated the adaptive value of priming within the same generation and showed that it is a distinct immune strategy that can evolve independently of basal resistance (without priming) to infection susceptibility in the flour beetle *Tribolium castaneum* (Khan et al., 2017a). Also, a striking result of this study was that the net survival benefit of evolved priming was considerably lower than that of resistance (50% vs 80% survival after infection; Khan et al., 2017a), suggesting their inherently different effects on overall infection outcomes. However, the selective forces influencing the evolution of these diverse insect immune strategies and infection outcomes remain largely unclear.

Do different evolutionary forces shape insect priming vs basal resistance to infection? The potentially divergent costs of maintenance and deployment of various immune responses can shed some light here (Norris and Evans, 2000; Sheldon and Verhulst, 1996). Several studies suggest that resistance is associated with overexpression of fast-acting immune responses that impose large physiological costs (e.g. Sadd and Siva-Jothy, 2006; Khan et al., 2017b). A general mathematical model predicts that such costs of constitutively expressed basal resistance can be outweighed by its benefit only under frequent lethal pathogenic infections, maximising the population’s growth rate (Mayer et al., 2016). However, the cost of resistance may be larger when pathogens are encountered infrequently. This is perhaps one reason why our beetle populations infected with a single large dose of infection every generation evolved priming (Khan et al., 2017a). In contrast, resistance could evolve only in populations that were exposed repeatedly to the pathogen within the same generation (primary exposure with heat-killed bacteria followed by live bacterial infection) (Khan et al., 2017a). However, few studies have actually measured the fitness consequences of evolved resistance in insects, and these were equivocal: e.g. while some found significant costs in terms of reduced longevity, poor larval competitive ability and delayed development (Kraaijeveld and Godfray, 1997; Ye et al., 2009; Ma et al., 2012), several others did not (Faria et al., 2015; Gupta et al., 2016). In general, costs of pathogen resistance may manifest as widespread tradeoffs with other life-history parameters as well, including reproduction (Reviewed in Sheldon and Verhulst, 1996; Lochmiller and Deerenberg, 2000; Rolff and Siva-Jothy, 2003; Schwenke et al., 2016). In contrast, the impact of immune priming-mediated protection on various fitness parameters has only recently been tested: e.g., primed mosquitoes (Contreras-Garduño et al., 2014), tobacco hornworms (Trauer and Hilker, 2013) and flour beetles (Khan et al., 2019) show reduced fecundity or primed mealworm beetle mothers produced progeny that developed slowly (Zanchi et al., 2011). Although these experiments revealed diverse fitness impacts of mounting priming responses using phenotypic manipulations, the actual costs of evolving, maintaining and deploying priming responses in populations while facing persistent pathogen-imposed selection have never been measured in detail (also see Ferro et al., 2019).

We also considered the potential added fitness benefits of evolved priming, where survival benefits manifest not only in the primed individuals (i.e., WGIP) but also across generations (i.e., TGIP). Besides enhancing the net fitness impact of priming via WGIP, TGIP might also facilitate the evolution and spread of priming ability in populations. Although no direct experiments have tested whether such trans-generational benefits evolve simultaneously with WGIP, theory offers some important clues. A model by Tidbury and coworkers suggests that since TGIP has a lower ability to reduce infection prevalence, selection should favour WGIP more (Tidbury et al., 2012). On the other hand, Tate and Rudolf suggested that the stage-specific effects of infection are important: TGIP is more beneficial when an infection affects juvenile stages, whereas WGIP is more effective if adults incur higher infection costs than larvae (Tate and Rudolf, 2012). The model also predicts that selection can strongly favour both WGIP and TGIP when the pathogen affects larvae and adults equally (Tate and Rudolf, 2012). Our previous experimental results suggest that this hypothesis might be relevant at least for flour beetles: both WGIP and TGIP were equally beneficial in beetles infected with the general insect pathogen *Bacillus thuringiensis* (Bt), which imposed similar infection costs across life stages (Khan et al., 2016). Although these results represent an interesting correlation, the causal link between the pathogen’s impact on the host and its role in determining relative investment in different priming responses is not yet confirmed.

To understand the selective pressures and fitness effects that directly impact the evolution of diverse priming responses vs basal infection resistance, we used previously described, evolved replicate populations of the red flour beetle *T. castaneum* that were infected in each generation with Bt (DSM 2046), either with or without the opportunity of priming with heat-killed Bt cells (described in Khan et al., 2017a; also see **Fig.** 1A and Supplementary methods). Earlier, we had analysed evolved infection responses of these populations after 11 generations of evolution (Khan et al., 2017a): (i) post-infection survival benefits of primed individuals (relative to their unprimed counterparts) as effects of immune priming, and (ii) the inherent ability to survive after infection (without prior priming with heat-killed cells) with respect to sham infected controls, as a proxy for basal resistance against infections. In the present study, we re-tested the same populations after a further 3 generations of experimental evolution to analyse the underlying fitness costs and benefits of their evolved responses. We measured a range of critical fitness-related traits in these populations such as offspring development, early reproduction and early survival, as well as adult lifespan under starvation and normal conditions. We also tested whether the evolution of priming ability was associated with transgenerational survival benefits during infection.

**Figure 1.**
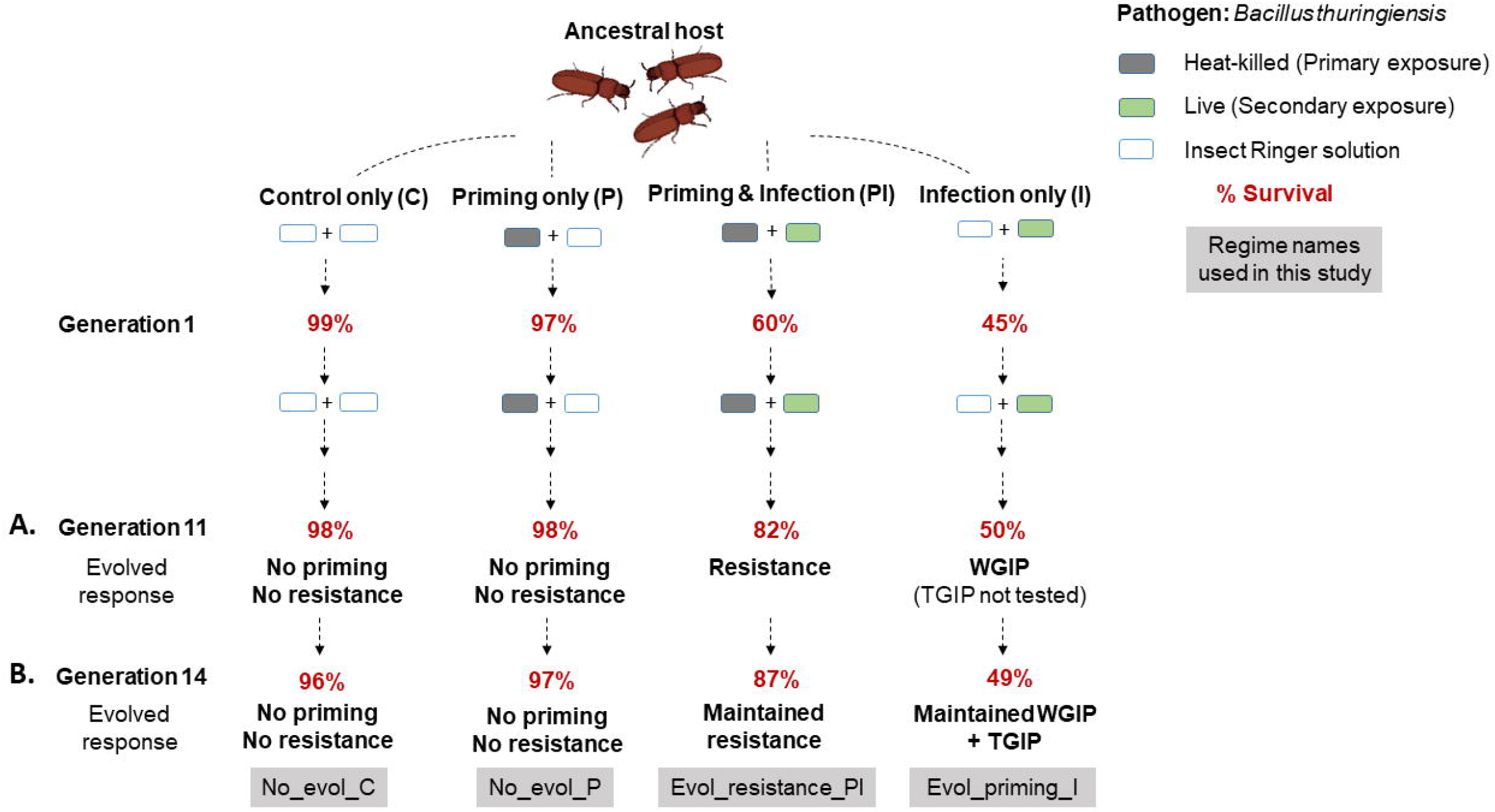
Summary of the design and outcome of experimental evolution of *Tribolium castaneum* flour beetles against the bacterial pathogen *Bacillus thuringiensis*. The schematic indicates beetle survival as well as evolved immune strategies (resistance or different priming forms) observed in all populations of each regime, after **(A)** 11 and **(B)** 14 generations of experimental evolution, described in Khan et al., 2017a and the present study respectively. C, P, PI and I populations assayed at 11^th^ generation (Khan et al., 2017a) were renamed as no_evol_C, no_evol_P, evol_resistance_PI and evol_priming_I populations respectively in the present study. Every generation, 10-day-old virgin beetles were either injected with heat-killed bacterial slurry (no_evol_P & evol_resistance_PI beetles) or sterile insect Ringer solution (no_evol_C & evol_priming_I beetles) (primary exposure) (See **SI** methods). After six days, individuals from evol_priming_I and evol_resistance_PI regimes were challenged with live Bt, whereas no_evol_C and no_evol_P beetles were pricked with sterile insect ringer solution (secondary exposure) (See **SI** methods). In the current study, we analysed 3 replicate populations from each regime. WGIP = Within-generational immune priming; TGIP = Trans-generational immune priming.

Note that we did not measure the direct changes in immune function or pathogen clearance in our experiments. Hence, the resistance, estimated from post-infection survival data alone, can be confounded by the potential evolution of tolerance, whereby survival might increase by reducing the cost of infection or immune responses, not because of more efficient pathogen killing mechanisms (Ayres and Schneider, 2012). Nonetheless, our work provides the first systematic analysis of the evolutionary cost and benefit structure influencing parallelly evolved, divergent survival responses to pathogens infections, by using possibly different immune strategies. We also provide the first report of the experimental evolution of TGIP in insects.

## 2. METHODS

### 2.1 Experimental evolution

We used laboratory-adapted populations of *T. castaneum* to initiate four distinct selection regimes, described in (Khan et al., 2017a): control (previously C; henceforth “no_evol_C”), priming only (previously P; henceforth “no_evol_P”), primed and infected (previously PI; henceforth “evol_resistance_PI”) and infection only (previously I; henceforth “evol_priming_I”), each with originally 4 independent replicate populations (also outlined in **Fig.** 1). In the present study, for logistical reasons, we could only analyze three replicates from each selection regime (no_evol_C 1, 2 & 4; no_evol_P 1, 2 & 4; evol_resistance_PI 1, 2 & 4; evol_priming_I 1, 2 & 4). On different days, we handled one replicate population from each selection regime together. The detailed protocol for the experimental evolution is described in (Khan et al., 2017a) (also see Supplementary methods), where we quantified evolved responses after 11 generations of selection (**Fig.** 1A). In the current study, we reanalyzed these evolved lines after three further generations of selection (**Fig.** 1B). Thus, after 14 generations of continuous selection, we isolated a subset of individuals from each replicate population and propagated them without any selection for two generations, i.e., without priming or infection (unhandled). This relaxing of selection is expected to generate standardised experimental beetles with minimum non-genetic parental effects.

### 2.2 Joint assays of evolved priming and resistance, and their impacts on reproduction

We designed our experimental framework to jointly compare survival benefits and reproductive effects of evolved priming vs. resistance against Bt infection (see **Fig.** 2A for experimental design). Besides measuring survival after priming and infection, we measured female reproductive output both before and after infection. This allowed us to estimate the direct impact of experimental evolution with pathogens vs. the actual impact of inducing each type of immune response. Simultaneously, we also tested for the evolution of trans-generational immune priming (TGIP), to compare relative survival and reproductive effects of different priming responses.

**Figure 2.**
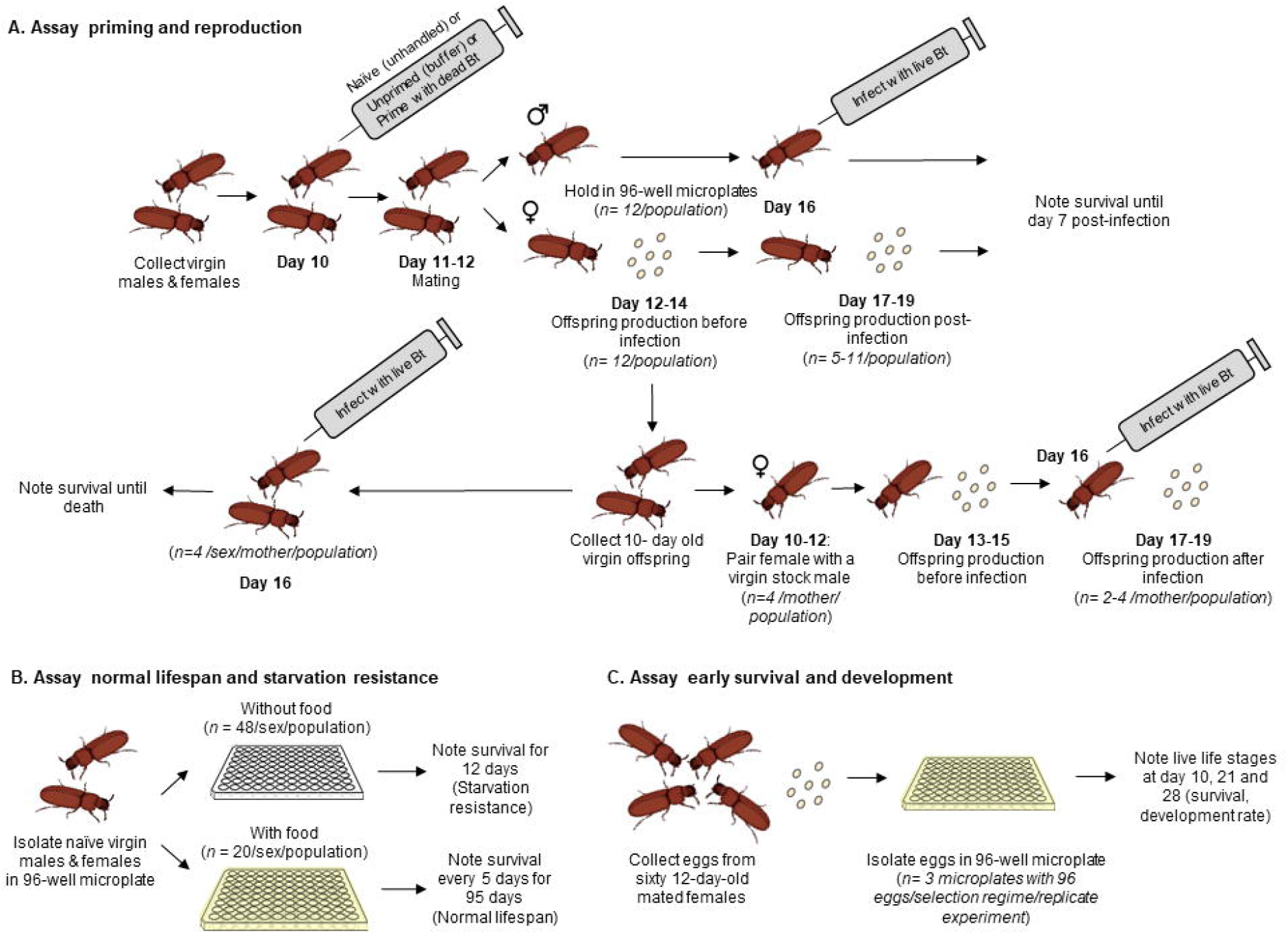
Design of experiments to assay **(A)** evolved immune responses and their impacts on beetle reproduction **(B)** survival under normal condition and starvation **(C)** early survival and development.

#### 2.2.1 Confirmation of evolved immune responses, and measurement of their reproductive effects (see **Fig.** 2A for experimental outline)

We first collected pupae from each standardised population and isolated them into 96-well microplate wells with ~0.2g wheat flour, for eclosion. We randomly assigned 10-day old virgin males and virgin females from each population to one of the following primary exposure treatments: (a) naïve (or unhandled) (b) primed (injected with heat-killed Bt) and (c) unprimed (i.e., injected with insect Ringer) (see **SI** methods for various priming and infection treatments). After 24 hours of primary exposure, we formed mating pairs using males and females within the same primary exposure treatment combination from each population in 1.5ml micro-centrifuge tubes with 1g of wheat flour (n = 12 mating-pairs/ priming treatment/ replicate population/ selection regime). We allowed them to mate for 48 hours and then isolated the 12-day-old females to oviposit for another 48 hours in 5 g whole wheat flour (oviposition plate), whereas males were returned to 96-well microplates. After oviposition, we also returned the 14-day-old females to 96-well microplates. Two days later (total six days after primary exposure), we infected males and females with live Bt. We recorded male survival every 6 hours for 1 day and then every 24 hours for 7 days post-infection (same as the selection window during experimental evolution; Khan et al., 2017a). We tracked female survival similarly, except that a day after infection, we again allowed 48-hour oviposition to estimate the reproductive impact of infection and induction of any priming responses. Here, we caution that since bacterial infection imposed significant mortality across regimes, the replicate size for our fitness assays was lower than expected (n=5-12 females/ priming treatment/ replicate population/ selection regime; **Table** S1; resulting analyses can thus be relatively underpowered compared to the reproductive assays performed before infection). We also note that although more beetles were alive in the evol_resistance_PI regime during the reproductive assay, we did not find a significant difference in the proportion of live beetles that reproduced across different treatments and selection regimes (**Table** S1). We also conducted a mock challenge for a subset of unprimed beetles as a procedural control for the survival assay, but not for reproductive output. We did not find any mortality in uninfected beetles within the experimental window of 7 days.

To test whether reproductive costs of evolved resistance manifest under stressful conditions, we compared the reproductive output of naïve unhandled evol_resistance_PI beetles with no_evol_C beetles reared in a poor-quality diet, corn (Agashe et al., 2011). To generate experimental adults, we allowed females from standardized no_evol_C and evol_resistance_PI populations (replicate populations 1 and 2) to oviposit in cornflour and collected their offspring to measure reproductive output as described above (n=36 females/replicate population/selection regime).

#### 2.2.2 Generation of beetles (progeny) for transgenerational priming (TGIP) assay

We used eggs laid by naïve, unprimed and primed females (assayed to estimate the effects of WGIP, described above) to also measure TGIP. We allowed the eggs to develop for 21 days at 34ºC and counted the total number of progeny (mostly pupae). Particularly, we retained the offspring from the first round of oviposition (without infection) (See **Fig.** 2A). At this time, most offspring were pupae, and the few adults we observed had pale body coloration indicating that they were not sexually mature and hence, unlikely to be mated (Sokoloff, 1977). We isolated these pupae and adults in 96-well plates with ~0.2g flour, to obtain virgin beetles for assays to measure trans-generational priming and offspring reproduction. We included offspring from mothers that produced more than 8 female and 4 male offspring (n= 8-10 mothers/ priming treatment / replicate population/ selection regime), enabling us to sample enough beetles to jointly test offspring post-infection survival (a proxy of trans-generational priming; described below) and reproduction of each parental pair.

#### 2.2.3 Assaying transgenerational priming response and its reproductive effects (See Fig. 2A for experimental outline)

After 10 days, we combined a subset of female offspring from each parental pair (n=4 offspring/ parental pair/ priming treatment/ replicate population/ selection regime) with 10-day old virgin males from standard laboratory stock population into a single mating pair for 48 hours and then allowed them to oviposit. This enabled us to measure the impact of parental priming on offspring reproductive output across populations. On day 16, we infected females and then again assayed their reproductive output as described above. We could only compare the post-infection reproduction of primed and unprimed females, but not naïve females as we lost them accidentally. On the same day, we also infected the remaining 16-day old virgin male and female offspring from each parental pair with live Bt (n= 4 offspring/ sex/ parental pair/ priming treatment/ replicate population/ selection regime) and noted their survival every 6 hours for 2 days and then every 24 hours until all of them were dead. This experimental design not only allowed us to jointly estimate the survival and reproductive effects of WGIP vs. TGIP for each parental pair, but also to analyse the impact of each type of immune response relative to evolved resistance. We did not find any mortality in sham infected offspring within the experimental window.

### 2.3 Quantifying development and survival under starvation and with food, in evolved lines

In separate experiments, we measured the direct impacts of evolved priming responses and resistance on other fitness components of naïve unhandled beetles (without priming and challenge), such as early survival, development, starvation resistance and lifespan.

#### 2.3.1 Impact on lifespan under starvation and with food (See Fig. 2B for experimental outline)

We first isolated 10-day old naïve virgin males and females from each population in 96-well microplate wells without food (n = 20 beetles/ sex/ replicate population/ selection regime). We noted mortality every 12 hours (10 am & 10 pm ±1 hour) for the next 12 days until all beetles died (**Fig.** 2B). In a separate experiment, we similarly distributed naïve virgin females into 96-well microplates, but with access to food (n = 48 females/ replicate population/ selection regime). We noted their survival every 5 days for 95 days to estimate the long-term survival costs of evolved immune responses. We could not assay males for long-term survival costs due to logistical challenges.

#### 2.3.2 Quantifying early survival, development and viability costs in evolved lines (See **Fig.** 2C for experimental outline)

We next estimated the impact of evolved immune responses on aspects of early survival and development. We allowed 12-day old, mated females from each population (n = 60/ replicate population/ selection regime) to oviposit in 150g of doubly sifted flour (using sieves with pore size of 50μ to remove large flour particles; Diager USA) for 24 hours. We had 3 replicates for each population. We discarded the females, and isolated 96 randomly chosen eggs into 96-well microplate wells with ~0.2 g flour (n= 3 microplates/ replicate population/ selection regime). This method was designed to detect intrinsic patterns of several developmental features across populations, without the confounding effects of density-dependent competition at the juvenile stage. After 10 days, we sifted the flour from each microplate to count live larvae and measure egg hatchability. Following this, we again returned the live larvae to 96-well plates and provided fresh flour. In our standard stock beetle populations, pupation and adult emergence begins around 3-4 weeks after oviposition. Therefore, we estimated the proportion of pupae and adults after 3- and 4-weeks post-egg collection respectively, as proxies for time to pupation and adult emergence. Finally, we calculated the percentage of surviving larvae, pupae and adults at week 4, as a proxy for overall viability. Since no_evol_P beetles did not evolve priming or resistance, we excluded them from the assay.

### 2.4 Data analysis

#### 2.4.1 Within generation priming response and resistance in parents

We first analysed survival data for all selection regimes using a mixed effects Cox model using the R package ‘coxme’ (Therneau, 2015). We fit separate models for females and males, since they were assayed on different days. In each case, we specified the model as survival ~ selection regime × priming treatment + (1|selection regime/ replicate population), with selection regime and priming treatment as fixed effects, and replicate populations nested within the selection regime as a random effect. A significant interaction between priming treatment and selection regime would indicate that the effects of priming varied across selection regimes. We note that although this analysis provides an overall estimate of each effect, complex interactions can prevent us from making meaningful comparisons between selection regimes. To understand the differences across priming treatments (i.e., unprimed vs primed) and selection regimes in detail, we therefore separately analysed survival data for each standardised replicate population, using Cox Proportional Hazard survival analysis. We noted individuals that were still alive at the end of the survival experiment as censored values and estimated the survival benefit of priming as the hazard ratio of unprimed versus primed groups. A hazard ratio significantly greater than one indicates a higher risk of mortality in the unprimed group relative to primed individuals; hence, a significant survival benefit of WGIP.

Separately, we also estimated the hazard ratio of naïve infected beetles from the no_evol_C regime against naïve infected beetles from all other regimes, to quantify evolved resistance against Bt infection. While a hazard ratio significantly greater than one indicates a higher risk of mortality in no_evol_C beetles, it also implies increased evolved resistance in selected regimes relative to no_evol_C beetles.

#### 2.4.2 Transgenerational priming response in offspring

To measure TGIP, we recorded the survival of 4 male and 4 female replicate offspring from each parental mating pair assayed earlier for within-generation priming. We calculated the mean lifespan for both the sexes as the unit of analysis but noted that residuals of lifespan data were not normally distributed (verified with Shapiro-Wilk tests; p<0.01). Therefore, we transformed the data into square root values that fit a normal distribution, and then compared group means of male and female offspring survival using a linear mixed effects model (R package ‘lme4’, Bates et al., 2012) with selection regime, parental priming status and offspring sex as fixed factors and replicate populations as a random effect. We tested for pairwise differences between selection regime and treatment after correcting for multiple comparisons using Tukey’s HSD (honest significant difference), using the R package ‘lsmeans’ (Lenth, 2017). We performed model reduction through stepwise removal of nonsignificant terms.

In addition to detecting the overall TGIP response in evol_priming_I beetles, we also wanted to compare their relative survival benefit with that of WGIP. To this end, for each replicate population, we first separately analysed the lifespan of male and female offspring from each parental pair, using a mixed effects Cox model to calculate the estimated hazard ratio of offspring from unprimed parents versus primed parents. Since we recorded the survival of 4 male and 4 female offspring from each parental pair, we specified the model as: survival ~ parental priming status + (1| parental mating pair/ replicate offspring), with parental priming status as a fixed effect and male (or female) replicate offspring nested within each parental mating pair as a random effect. Since all beetles died within the experimental window, we did not have any censored beetles in the analysis.

Subsequently, we used non-parametric Wilcoxon Rank Sum tests to compare hazard ratios calculated from TGIP versus WGIP for each population.

#### 2.4.3 Reproductive effects

We found that the residuals of pre-infection reproductive output data of both parents and offspring were non-normally distributed and could not be transformed to a normal distribution. We, therefore, used non-parametric Wilcoxon Rank Sum tests to analyse the impact of selection regime and priming treatment (for replicate populations of no_evol_C, no_evol_P, evol_resistance_PI and evol_priming_I that were handled together). We also used Wilcoxon tests to analyse the impact of bacterial infection on the reproductive output of parents and offspring, separately for each replicate population across selection regimes and priming treatments.

We analysed post-infection reproductive fitness data using a linear mixed effects model with selection regime and priming treatment as fixed factors and replicate populations as a random effect [Model: No. of offspring~ Priming treatment × Selection regime + (1 ⍰ Replicate population); fitted separately for parents and offspring]. To understand the complex interactions between treatment and selection regime and to disentangle the effects of each type of evolved immune response (i.e., priming in evol_priming_I or resistance in evol_resistance_PI), we also compared post-infection reproductive data of parents from each selection regime separately with that of control beetles (no_evol_C), wherever needed. We used Tukey’s HSD to test for pairwise differences between selection regimes and treatments. As described above, we removed the nonsignificant effects stepwise to determine the best reduced model.

#### 2.4.4 Survival under starvation and fed conditions

We analysed survival data under starvation and with food, using mixed effects Cox models as described below (best reduced model, wherever needed):

I. Lifespan under starvation: survival ~ selection regime + sex + (1| selection regime/ replicate population), with selection regime and sex as fixed effects, and replicate populations nested within selection regimes as a random effect.
II. Lifespan with food: survival ~ selection regime + (1| selection regime/ replicate population), with selection regime as a fixed effect, and replicate populations nested within selection regimes as a random effect.

#### 2.4.5 Early survival, development and viability costs

We analysed data using a linear mixed effects model with selection regime as fixed factor and replicate population as a random effect. In each case, we described the model as: Trait ~ Selection regime + (1| Replicate population). We tested for pairwise differences using Tukey’s HSD.

## 3. Results

Our previous work demonstrated that lethal Bt infection rapidly selects for divergent post-infection survival responses (potentially via divergent immune strategies in beetles), within 11 generations (Khan et al., 2017a). Populations that were directly infected with a single large dose of Bt evolved within-generation priming response (evol_priming_I regime), whereas populations, where beetles were injected first with heat-killed and then live Bt, evolved basal resistance to infection (evol_resistance_PI) (**Fig.** 1A). Our results after 14 generations of selection were consistent with these prior observations (Khan et al., 2017a) (**Fig.** 1B). A mixed-effects Cox model fitted separately to male and female survival data revealed a significant interaction between selection regime and treatment, suggesting that effects of priming and infection varied across selection regimes (**Fig.** S1-2, **Table** S2-3).

To understand these complex effects in detail, we next, analysed survival data for each population with a Cox proportional hazard analysis. We found evolved priming responses only in males and females from evol_priming_I populations (~3-fold increase in their survival relative to control beetles) (**Fig.** S1-2, **Table** S2-S3); whereas evol_resistance_PI beetles had higher basal resistance (3 to 28-fold increase in the survival of naïve evol_resistance_PI beetles relative to control beetles) (**Fig.** S1-2, **Table** S4). We also found that while the survival of evol_priming_I beetles after Bt infection was still 50%, evol_resistance_PI beetle survival had increased to ~85% (**Fig.** S3). As expected, populations from the no_evol_C or no_evol_P regimes (where beetles were not exposed to live infection) did not evolve any priming ability or higher resistance to infection.

### 3.1 Maintenance of evolved immune responses do not incur reproductive costs

We measured the impact of evolved immune responses on beetle reproduction and found complex fitness effects that varied substantially with priming type and infection status. Evolved priming or resistance had no significant impact on the reproduction (**Fig.** 3, **Table** S5). There were also no general effects of priming treatment on beetle reproduction (**Table** S5). Thus, the maintenance of evolved priming or resistance does not impose a reproductive cost. However, we noticed that females infected with live pathogen later in their life (see methods) had reduced reproductive output in most populations, except those from evol_resistance_PI regime where the effect was relatively mild (compare across selection regimes and treatments; **Fig.** 3, **Table** S6). Thus, most evol_resistance_PI beetles, regardless of their priming status, generally did not exhibit reproductive costs of infection. Also, the average post-infection reproductive cost of evolved priming appeared higher than that of evolved resistance (compare beetles from evol_resistance_PI vs evol_priming_I populations in **Fig.** 3, **Table** S6), highlighting contrasting outcomes of live infection across evolved regimes. These patterns were further confirmed in subsequent analyses where we separately compared the reproductive fitness of naïve and unprimed females after infection, using a Wilcoxon rank sum test (data was not normally distributed; Shapiro-Wilk test: P<0.05). We did not include primed beetles in these analyses, because we wanted to measure the effect of live infection without any interference from previous antigen exposure (i.e., heat-killed bacteria during priming). Overall, there was no effect of the experimental treatment (naïve vs unprimed females after infection: df=1, χ2= 0.021, P=0.87), but overall infected evol_resistance_PI beetles produced significantly more offspring than no_evol_C (df=3, χ2= 12.458, P<0.001) and evol_priming_I beetles (df=3, χ2= 14.837, P<0.001) beetles.

**Figure 3.**
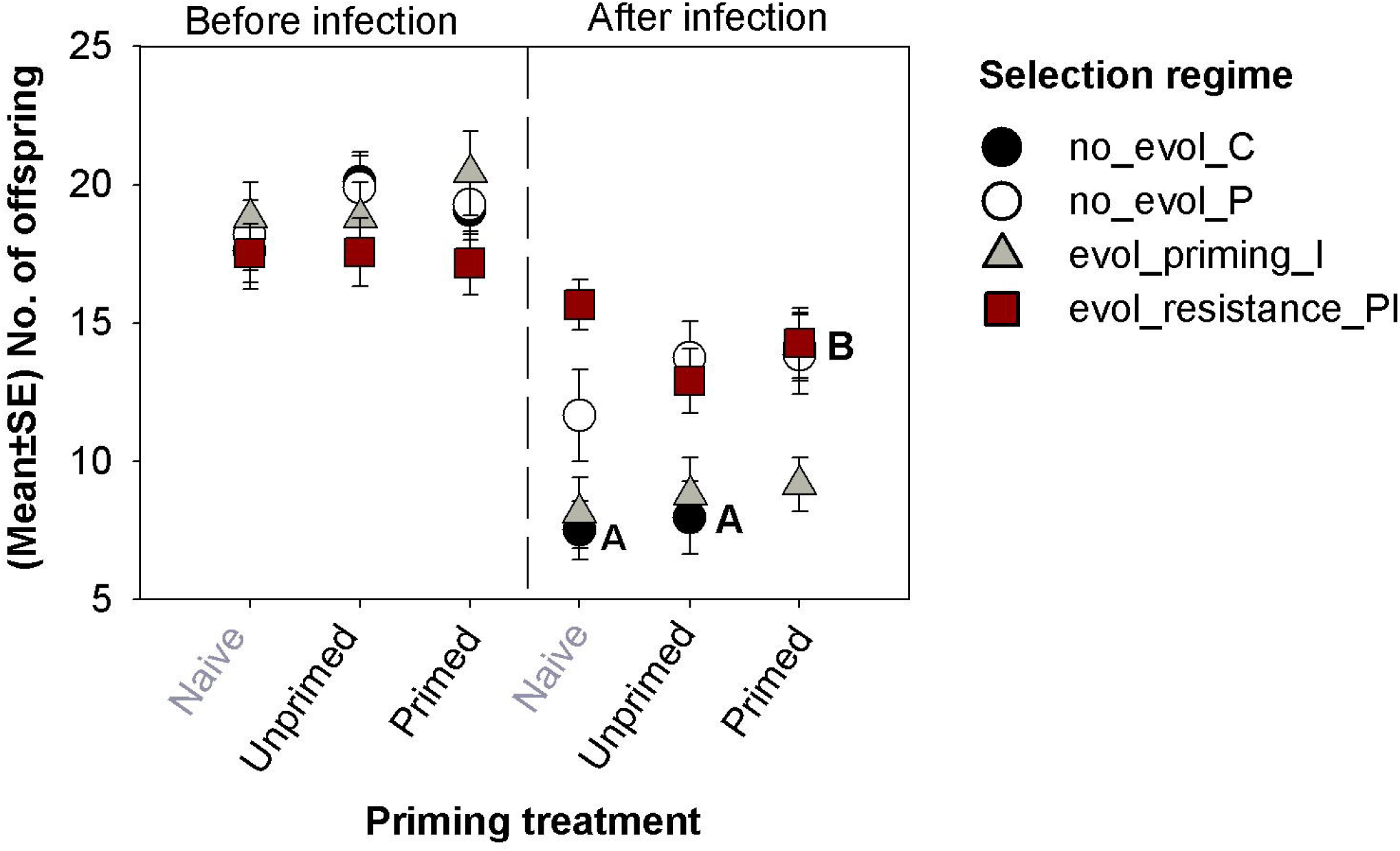
Impact of evolved within-generation priming (WGIP) and resistance on female reproductive output, both before (n=12 females/ priming treatment/ replicate population/ selection regime) and after bacterial infection (n=5-11 females/ priming treatment/ replicate population/ selection regime) (partitioned by the vertical line). Different letters denote significant differences in post-infection reproduction of no_evol_C beetles after mounting a within-generation priming response (tested with Tukey’s HSD). Error bars denote standard errors.

Next, we analysed the full dataset (including naïve, unprimed & primed females) to study the impact of experimental priming (mimicking selection regimes during experimental evolution) on the reproductive output of infected beetles. We expected that after infection, beetles in the priming treatment would reflect reproduction of evol_resistance_PI beetles; whereas beetles in the unprimed treatment would mimic evol_priming_I beetles during experimental evolution. In comparison to no_evol_C beetles, they would thus provide an estimate of the reproductive effects of mounting evolved resistance and priming response respectively during actual infection. A linear mixed effects model with reproductive output data from all regimes revealed significant main effects of both selection regime and treatment as well as their interaction (**Table** S7 I). Beetles from the evol_resistance_PI (P=0.04) and no_evol_P (P=0.02) regimes appeared to produce more offspring than no_evol_C beetles, though these effects were perhaps confounded by their priming status. Since our experiments were primarily designed to disentangle the reproductive effects of separately evolved responses with respect to control no_evol_C beetles, we also compared each selection regime individually with no_evol_C control beetles (e.g., evol_priming_I vs no_evol_C, or evol_resistance_PI, vs no_evol_C) (**Table** S7 II-IV). In contrast to results obtained from the full dataset (**Table** S7 I), these comparisons did not show consistent effects of the selection regime, but revealed significant effects of the priming treatment and its interaction with selection regime in each case (**Table** S7 III-IV). In both analyses (i.e., full dataset and pairwise comparisons), experimental priming induced a large increase in reproduction only in no_evol_C regime after infection (primed vs unprimed no_evol_C; Tukey’s HSD: P=0.005), but not in beetles from other selection regimes (Tukey’s HSD: p>0.9) (**Fig.** 3). Surprisingly, primed evol_priming_I beetles failed to increase their reproduction despite increased post-infection survival benefits (after mounting WGIP), suggesting a fitness cost. This was in contrast to evol_resistance_PI lines that could apparently alleviate this reproductive cost by maintaining higher reproduction than no_evol_C beetles regardless of their priming or infection status (discussed above). Thus, evolved resistance is better than priming not only in terms of its survival benefit, but also in terms of reproduction. We also verified these significant differences by calculating confidence intervals (2.5%, 97.5%) [no_evol_C: naïve (4.633, 10.366), unprimed (5.528, 10.391), primed (11.595, 16.665) vs. evol_ priming_I: naïve (5.595, 10.665), unprimed (6.108, 11.415), primed (6.977, 11.345) vs. evol_resistance_PI: naïve (13.269, 18.038), unprimed (10.719, 15.087), primed (12.046, 16.486)].

Finally, in a separate experiment, we demonstrated that evol_resistance_PI beetles reared in poor quality diet corn produced an equal number of offspring to that of no_evol_C beetles (**Fig.** S4; **Table** S8: p=0.34), indicating that evolved resistance might not impose reproductive costs even under stressful conditions.

### 3.2 Evolved resistance increases survival under both starvation and fed conditions

We analysed the direct impacts of evolved priming and resistance on survival under starvation or fed conditions, using a mixed effects Cox model. In both cases, survival was significantly higher in evol_resistant_PI beetles compared to no_evol_C beetles (P<0.04 in each assay), but not in evol_priming_I or no_evol_P beetles (P>0.05 in each assay) (**Fig.** S5-6; **Table** S9-10). Males and females from all selection regimes had similar lifespan under starvation (**Fig.** S5, **Table** S9), suggesting no sex-specificity in starvation resistance.

### 3.3 Evolved priming reduces early survival and extends the development time

We measured features of early survival such as egg hatchability and the total number of viable offspring (i.e., live larvae, pupae and adults) within the first 4 weeks after oviposition. We also measured the proportion of pupae and adults at week 3 and 4, as proxies of development rate. We found significant effects of selection regime on the egg hatchability, the total number of viable offspring and proportion of adult offspring at week 4, but not on the proportion of pupae at week 3 (**Fig.** 4). The number of viable offspring at week 4 was drastically reduced in beetles from the evol_priming_I regime (**Fig.** 4D, **Table** S11). This was perhaps due to significant early mortality during egg to larval development in evol_priming_I beetles: while ~75% no_evol_C, no_evol_P and evol_resistance_PI eggs hatched into larvae, only 55% evol_priming_I eggs survived (**Fig.** 4A, **Table** S11). Besides, the proportion of adults at week 4 was lowest in the evol_priming_I regime, suggesting delayed development (**Fig.** 4C, **Table** S11). Overall, these results indicate that maintenance of priming imposed considerable costs in terms of reduced early survival and slower development in evol_priming_I beetles. In contrast, evolved basal resistance did not appear to impose a significant cost with respect to these traits.

**Figure 4.**
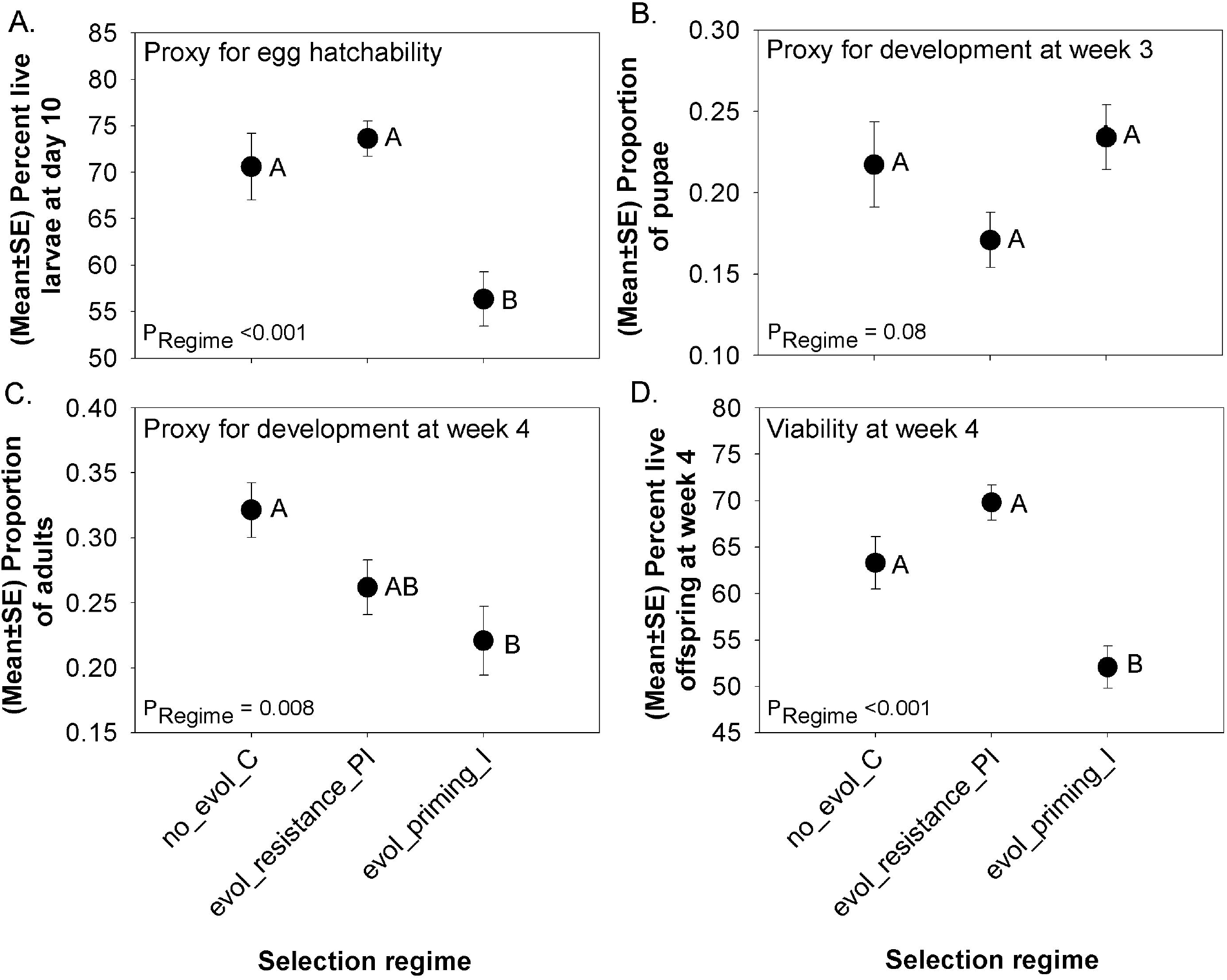
Impact of evolved immune responses on **(A)** total number of eggs that hatched into larvae (egg hatchability); proportion of **(B)** pupae at week 3 and **(C)** adults at week 4 as proxies for developmental rate; (D) the total number of viable offspring (larvae, pupae and adults) at week 4 (n= 3 × 96 well microplates/ replicate population/ selection regime). P values for the impact of the selection regime are reported in each panel. Significantly different selection regimes are indicated by distinct letters (based on Tukey’s HSD). Error bars denote standard errors.

### 3.4 Evolved WGIP is also associated with TGIP

Finally, we asked whether evolved priming conferred added trans-generational benefits, increasing its overall fitness impacts. We used a linear mixed effects model to analyse the mean post-infection survival of offspring from beetles assayed above as a function of selection regime, parental priming status and offspring sex. The selection regime and parental priming status had significant main effects as well as an interaction effect, whereas offspring sex had no impact (P>0.05) (**Fig.** 5A, **Table** S12). Here too, we found that overall, offspring of evol_resistance_PI beetles had the highest survival (compared with no_evol_C beetles across parental priming status, all comparisons with Tukey’s HSD: p<0.001), though they did not show effects of parental priming. In contrast, parental priming increased offspring survival in the evol_priming_I regime (Tukey’s HSD: p<0.001), suggesting that TGIP benefits are restricted to evol_priming_I beetles.

**Figure 5.**
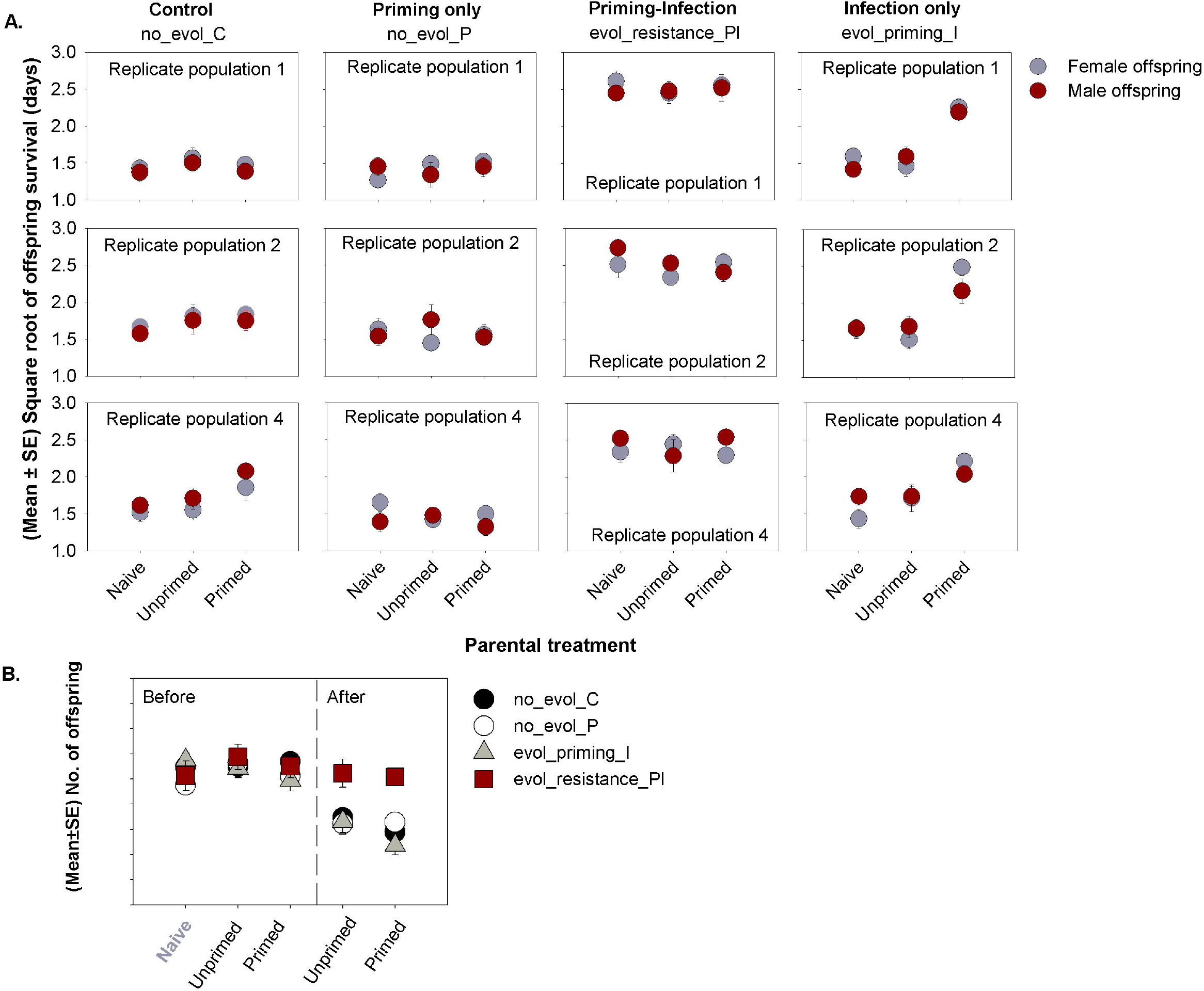
**(A)** Offspring survival after trans-generational immune priming (TGIP) and infection (group mean survival of 4 offspring from 8-11 parental pairs/ offspring sex / parental priming status/ replicate population/ selection regime). **(B)** Impact of evolved trans-generational priming on offspring’s reproductive output, both with and without infection (partitioned by the vertical line; group mean reproduction of 4 female offspring from 8-11 parental pair/ parental priming status/ replicate population/ selection regime). Error bars denote standard errors.

Separately, we also tested whether the relative survival benefits of TGIP were equal to that of WGIP. For each replicate population of evol_priming_I beetles and offspring sex, we thus used a mixed effects Cox model to analyze the individual offspring survival data nested within each parental mating pair, and calculated the strength of evolved TGIP as the estimated hazard ratio for offspring from unprimed vs. primed parents. We found a significant TGIP response in offspring from all replicate populations, except for male offspring from evol_priming_I4 parents (**Fig.** S7A, **Table** S13). Primed evol_priming_I4 male offspring appeared to survive more than their unprimed counterparts, but the difference in their survival was not statistically significant (**Fig.** S7A, **Table** S13). Nonetheless, the net survival benefit of TGIP and WGIP was not different across replicate populations (Wilcoxon Rank Sum Test: p>0.05), possibly supporting the hypothesis that Bt-imposed selection favours the evolution of both types of priming to a similar extent (**Fig.** S7B).

### 3.5 Evolved trans-generational priming (TGIP) does not impose a separate reproductive cost

As found with mothers (above), evolved priming and resistance did not consistently affect the reproductive output of naïve or uninfected offspring (**Fig.** 5B, **Table** S14); but infection generally reduced offspring reproductive output in all selection regimes except evol_resistance_PI (**Table** S15). A linear mixed effects model revealed significant main effects of selection regime and parental priming status (**Table** S16). Evol_resistance_PI beetles produced more offspring than control beetles (no_evol_C vs. evol_resistance_PI: P=0.009) (**Fig.** 5B); but the reproductive output of no_evol_P and evol_priming_I beetles did not differ from control (all P>0.05, Tukey’s HSD). Interestingly, parental priming reduced reproductive output in all beetles across selection regimes. However, since there was no significant interaction effect between selection regime and parental priming, we inferred that transgenerational survival benefits in evol_priming_I beetles did not impose any added reproductive costs, relative to no_evol_C beetles (**Fig.** 5B).

## 4. DISCUSSION

Previously, we showed that priming response and basal resistance against *Bt* infection evolve as divergent mutually exclusive strategies in flour beetles (Khan et al., 2017a). However, it was unclear if selective parameters underlying their evolution, and potential trade-offs, were also different. Here, we revisited these beetle lines to identify whether they actually incurred divergent fitness costs. Concomitantly, we also asked whether priming confers additional, trans-generational fitness benefits. To our surprise, we did not find any evidence for a cost of evolved resistance: it did not impact development and reproduction, and even increased survival during both starvation and normal conditions, contradicting the traditional view of immunity-fitness trade-offs (Ye et al., 2009; Ma et al., 2012). Instead, our data add to the growing body of work that suggests only a weak role for life-history trade-offs during the evolution of pathogen resistance (Faria et al., 2015; Gupta et al., 2016). Contrary to our expectation, evolved priming imposed diverse fitness costs such as reduced offspring early survival, development rate and reproduction. Interestingly, we found that priming ability was also associated with trans-generational benefits, representing the first evidence of experimentally evolved TGIP. However, the combined benefit of these two forms of priming (~50% survival after Bt infection) was still lower than that conferred by increased basal resistance to Bt (~85% survival).

Interestingly, we found that although infection reduced reproduction in all regimes, the effect was less pronounced in evol_resistance_PI beetles; hence, evolved basal resistance was also associated with a relative reproductive advantage. No_evol_P (priming only) beetles also had higher reproduction than control beetles after infection, which is counterintuitive because these beetles never experienced live infection during experimental evolution. Note that this relative reproductive advantage would be important during experimental evolution, since beetles reproduced for 5 days after infection in each generation (see methods). How do we interpret these apparent reproductive fitness benefits in evol_resistance_PI and no_evol_P beetles? First, as described earlier, the reduced cost of infection might also represent evolved tolerance (Ayres and Schneider, 2012), whereby beetles do not invest in directly clearing pathogens via canonical resistance mechanisms and may therefore be able to make a greater reproductive investment during infection. Since we could not compare the relative importance of tolerance vs resistance (e.g., by quantifying the fitness changes with increasing pathogen burden; Ayres and Schneider, 2012) in our beetle lines, future studies are necessary to test these possibilities explicitly. Second, our results could reflect a trade-off between early vs late reproduction. In other words, higher reproduction might represent a terminal investment in no_evol_P and evol_resistance_PI populations, whereas no_evol_C and evol_priming_I populations instead suppress immediate reproduction after infection to maintain survival and somatic maintenance later in life (Luu and Tate, 2017). However, long-term reproductive effects are probably not meaningful in the present context, because our beetles were selected to reproduce for only 7 days after infection. Overall, we suggest that divergent immune strategies can have important consequences for reproductive success, which deserve further attention.

We explored most of the plausible contexts in which immunity-fitness trade-offs could be expressed in our populations, given the experimental evolution regime wherein beetles evolved the divergent immune functions. It is possible that any underlying fitness trade-offs remained hidden because evol_resistance_PI beetles lived in a relatively stable environment, with *ad libitum* food (McKean et al., 2008). In contrast, suboptimal living conditions such as limited access to food and nutrition (McKean et al., 2008) could reveal fitness costs. To test this possibility, we compared the reproductive output of beetles from evol_resistance_PI lines vs. no_evol_C control lines when reared in a poor-quality corn diet (Agashe et al., 2011). Surprisingly, beetles that had evolved resistance to Bt still produced as many offspring as control beetles, indicating that evolved resistance did not impose reproductive costs even under stressful conditions. However, we want to reiterate that even if fitness costs of evolved resistance are expressed under other externally imposed environmental stresses (which could not be assayed in the present study), these conditions are not relevant for our selected populations that evolved under standard laboratory conditions. As such, any tradeoffs observed in these conditions cannot possibly explain the particular divergence in immune responses that we observed in our present experiment.

Our results might also contradict our prior hypothesis that at a low pathogen frequency (experienced by evol_priming_I beetles), priming may be more favourable than resistance due to its low maintenance costs (Khan et al., 2017a). Instead, we found that the overall maintenance of priming responses is costly. Although evolved priming did not affect lifespan or survival under starvation, it directly reduced egg hatchability, offspring viability and development rate in naïve evol_priming_I beetles compared to control beetles. These results broadly corroborate results from another beetle study showing that Bt-driven evolution of priming in larvae delayed their development (Ferro et al., 2019). However, such responses were observed only after mounting a priming response, suggesting a deployment cost; whereas basal maintenance costs in naïve beetles were not analysed. In our study, priming also had variable effects on reproduction across selection regimes. For instance, mounting a within-generation priming response helped no_evol_C beetles to increase their reproduction after infection; whereas infected evol_priming_I beetles, despite evolving survival benefits, could not improve their reproduction. These results mirror our recent observations with wild-caught populations, where primed and infected females with increased post-infection lifespan produced fewer offspring (Khan et al., 2019) and vice versa. We thus speculate that a hidden trade-off with post-infection reproduction might constrain the survival benefits of within-generation priming responses at a much lower level than resistance (see **Fig.** 1B). We also speculate that the evolution of costly priming perhaps requires reactive and self-damaging immune responses (e.g., phenoloxidase; see Ferro et al., 2017) to clear pathogens (Khan et al., 2017b; Khan et al., 2019) and hence physiologically constrains the extent of post-infection fitness benefits. In contrast, evol_resistance_PI beetles either used less toxic immune responses to clear pathogens or evolved tolerance, reducing immunopathological costs and maximising fitness benefit (Khan et al., 2017b). Mounting trans-generational priming responses, on the other hand, did not affect offspring reproduction, suggesting that fitness effects are not uniform across different types of evolved priming responses. Nevertheless, these results broadly corroborate other work showing the negative effects of priming on various fitness parameters (Trauer and Hilker, 2013; Contreras-Garduño et al., 2014). However, these studies primarily used phenotypic manipulations within a single generation, whereas ours is the first detailed study to directly measure the complex fitness costs associated with evolved priming across multiple generations of pathogen exposure.

Our experiments also provide the first empirical evidence that insects can evolve multiple priming responses simultaneously. Interestingly, both trans-generational and within-generation priming provided almost equivalent fitness benefits, corroborating our prior work showing similar benefits of WGIP and TGIP across 10 distinct wild-caught beetle populations (Khan et al., 2016). Such parallel results from natural and laboratory-evolved populations might indicate that natural pathogens such as Bt may serve as a potent source of selection favouring the evolution of diverse immune responses in insects. As discussed earlier, Bt reduces the survival of flour beetle larvae and adults equally (Khan et al., 2016), which should favour the simultaneous evolution of WGIP and TGIP (Tate and Rudolf, 2012). However, during experimental evolution, we only infected adult beetles, which should have restricted host-pathogen interaction to adults. It is possible that infected adults directly transmitted Bt to eggs, imposing selection favouring TGIP (see Tate and Rudolf, 2012). Alternatively, infected adults could have transmitted Bt (or antigen) to larvae via the flour, either through infected beetle cadavers (~10-15% mortality during oviposition period in evol_priming_I beetles) or excreta (Argôlo-filho and Loguercio, 2014). Another possibility is that ancestral beetle populations may have already coevolved with Bt in their natural habitat before they were brought into the lab. Consequently, despite being infected only as adults during experimental evolution, the beetle immune system could perhaps readily recognise Bt as a risk across life stages. Finally, if WGIP and TGIP involve shared molecular pathways, direct pathogen pressure on adults could result in simultaneous evolution of both types of priming. Although the molecular details responsible for immune priming are still unclear (Cooper and Eleftherianos, 2017) and perhaps confounded by diverse processes such as metabolic changes (Ferro et al., 2019) and epigenetic reprogramming of immune cells (Tate et al., 2017), recent data hint at shared immune pathways between different priming types. For instance, both within- (Pham et al., 2007) and trans-generationally primed bumble bees (Barribeau et al., 2016) show increased expression of Toll signalling pathways (also see Ferro et al., 2019). Further experiments to determine the molecules underlying different immune responses can help distinguish between the above hypotheses.

## 6. CONCLUSION

In closing, we note that the relative importance of priming vs general resistance has long been debated, primarily because it was unclear whether (a) diverse priming types (within-vs trans-generational) together constitute distinct strategies, separate from basal resistance (b) their costs vs benefits differ substantially, and (c) they involve different or overlapping sets of immune pathways. Our work represents one of the first steps to address the first two problems, demonstrating distinct costs and benefits of multiple priming responses vs resistance evolving simultaneously in response to selection imposed by the same pathogen. While these results highlight the remarkable diversity and flexibility of insect innate immune adaptation against infections, they also suggest that the early survival vs reproductive costs of priming can constrain its evolution, much more so than resistance. We hope that our results will motivate further experiments to understand this intriguing problem. Specifically, we look forward to detailed mechanistic studies to test how host-pathogen interactions at a low frequency of infection only favour the evolution of priming, mechanistically precluding more beneficial resistance alleles from fixing in host populations.

## ACKNOWLEDGEMENTS

We are grateful to Devshuvam Banerjee, Laasya Samhita, Srijan Seal and Shyamsunder Buddh for feedback on the manuscript. We thank Kunal Ankola, Sunidhi Thakur and Shyamsunder Buddh for their help during experiments, and Shivani Krishna and Basabi Bagchi for their help during the analysis.

## AUTHOR CONTRIBUTIONS

IK conceived experiments; IK, AP and DA designed experiments; AP carried out experiments; IK analysed data; IK and DA acquired funding; IK and DA wrote the manuscript with inputs from AP. All authors gave final approval for publication.

