## Supplementary Methods, Figure & Tables for "THE COSTS AND BENEFITS OF BASAL INFECTION RESISTANCE VS DIVERSE IMMUNE PRIMING RESPONSES IN AN INSECT"

**Priming and infection protocol**

We used a strain of *Bacillus thuringiensis* (DSM 2046) (Bt) originally isolated from a Mediterranean flour moth (Roth et al., 2009) as a model pathogen to prime and infect adult *Tribolium castaneum* beetles as outlined in Khan et al., (2017a). We used heat-killed bacteria to prime the immune system by activating the immune response without any direct cost of infection. Briefly, we first primed beetles by pricking them with a 0.1mm insect pin (Fine Science Tools, CA) dipped in heat-killed bacterial suspension prepared from freshly grown overnight Bt culture at 30°C (optical density of 0.95), or insect Ringer solution (mock priming) between their head and thorax. After six days, we infected individuals with a live and dense bacterial culture adjusted to ~10^10^ cells in 75 µl insect Ringer solution or pricked with sterile Ringer solution (mock challenge).

**Experimental evolution protocol**

Briefly, every generation, we first primed 10-day-old virgin no_evol_P and evol_resistance_PI adults from each replicate population with heat-killed bacterial slurry, as described above. Simultaneously, we also pricked virgin no_evol_C and evol_priming_I beetles with sterile insect Ringer solution (mock priming). Six days later, we challenged individuals from evol_priming_I and evol_resistance_PI regimes with live Bt, whereas no_evol_C and no_evol_P beetles were pricked with sterile insect ringer solution (mock challenge). We thus created two infection regimes where populations were challenged with a high dose of infection, with (evol_resistance_PI) or without (evol_priming_I) the opportunity of priming; and two control regimes where beetles were either pricked with Ringer (no_evol_C) or heat-killed bacteria (no_evol_P), but never exposed to live infection. Following the priming and infection treatments, we randomly isolated 60 pairs of live virgin males and females from each replicate population and provided them with 300g wheat to mate and oviposit for 5 days to initiate the next generation.

**SUPPLEMENTARY FIGURES**

**Figure S1.** Survival curves for within-generation priming and resistance in females (n= 12 males/ priming treatment/ replicate population/ selection regime) after 14 generations of selection. Asterisks and the numbers in parentheses for evol_priming_I beetles denote the hazard ratios calculated from survival curves for priming that are significantly greater than 1 (p<0.05; a greater hazard ratio indicates higher benefit of priming). Replicate populations of no_evol_C (C1, 2, 4), no_evol_P (P1, 2, 4), evol_resistance_PI (PI1, 2, 4) and evol_priming_I (I1, 2, 4) have been mentioned in each panel.

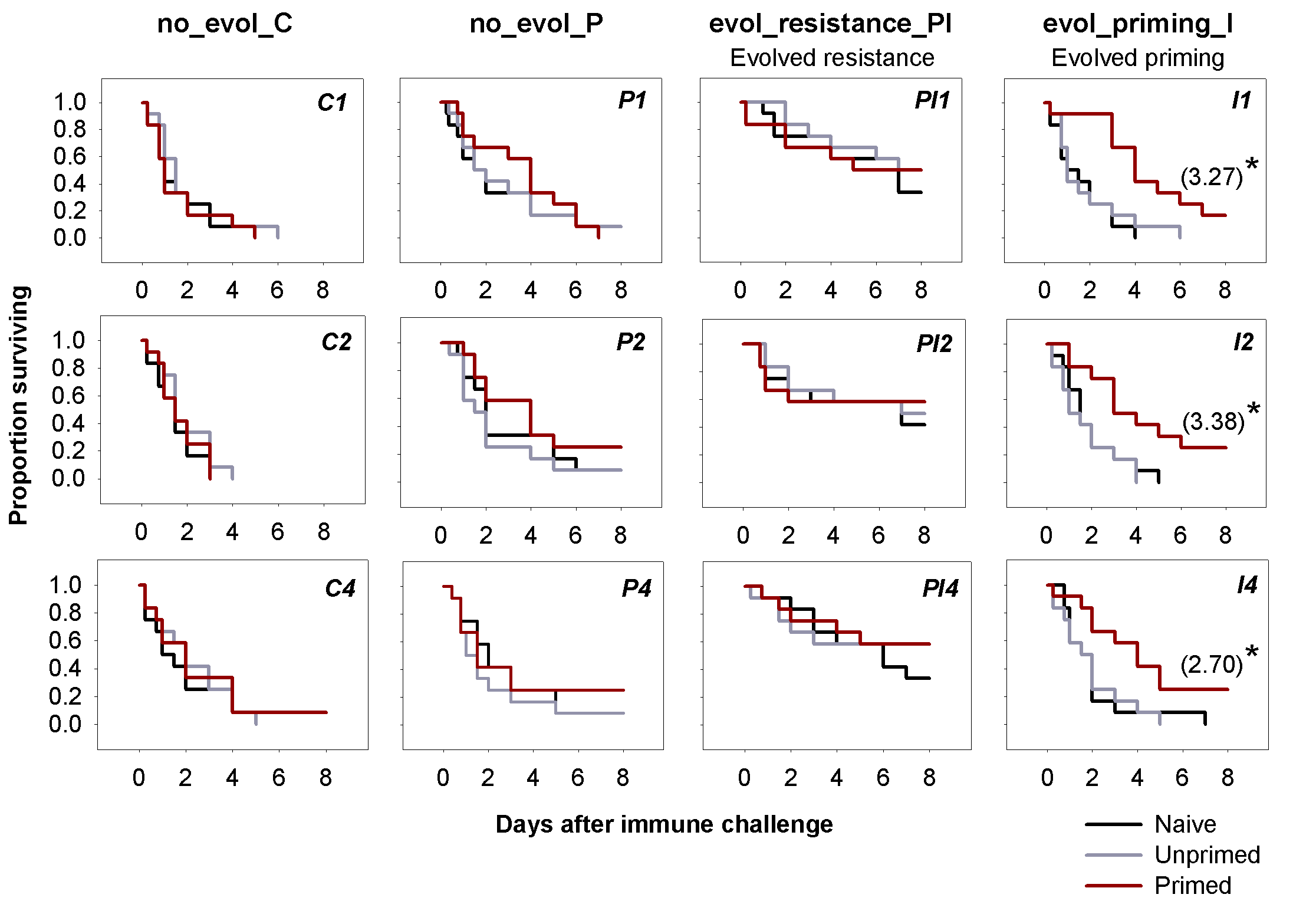

**Figure S2.** Survival curves for within-generation priming and resistance in males (n= 12 males/treatment/ replicate population/ selection regime) after 14 generations of selection. Asterisks and the numbers in parentheses for evol_priming_I beetles denote the hazard ratios calculated from survival curves for priming that are significantly greater than 1 (p<0.05; a greater hazard ratio indicates higher benefit of priming). Replicate populations of no_evol_C (C1, 2, 4), no_evol_P (P1, 2, 4), evol_resistance_PI (PI1, 2, 4) and evol_priming_I (I1, 2, 4) have been mentioned in each panel.

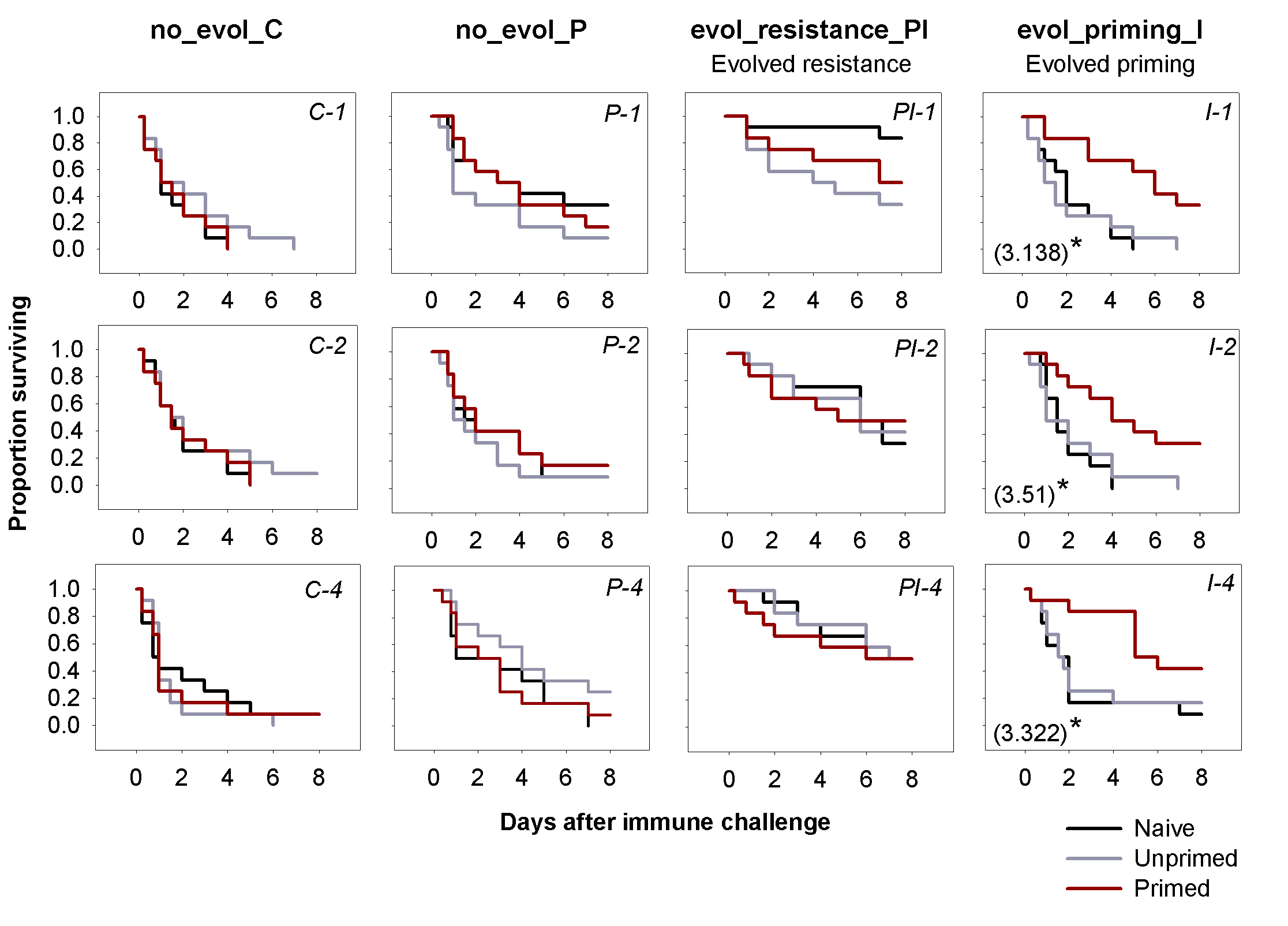

**Figure S3.** Adult survival after first 48h of infection with live Bt cells, during experimental evolution between generation 12 and 14.

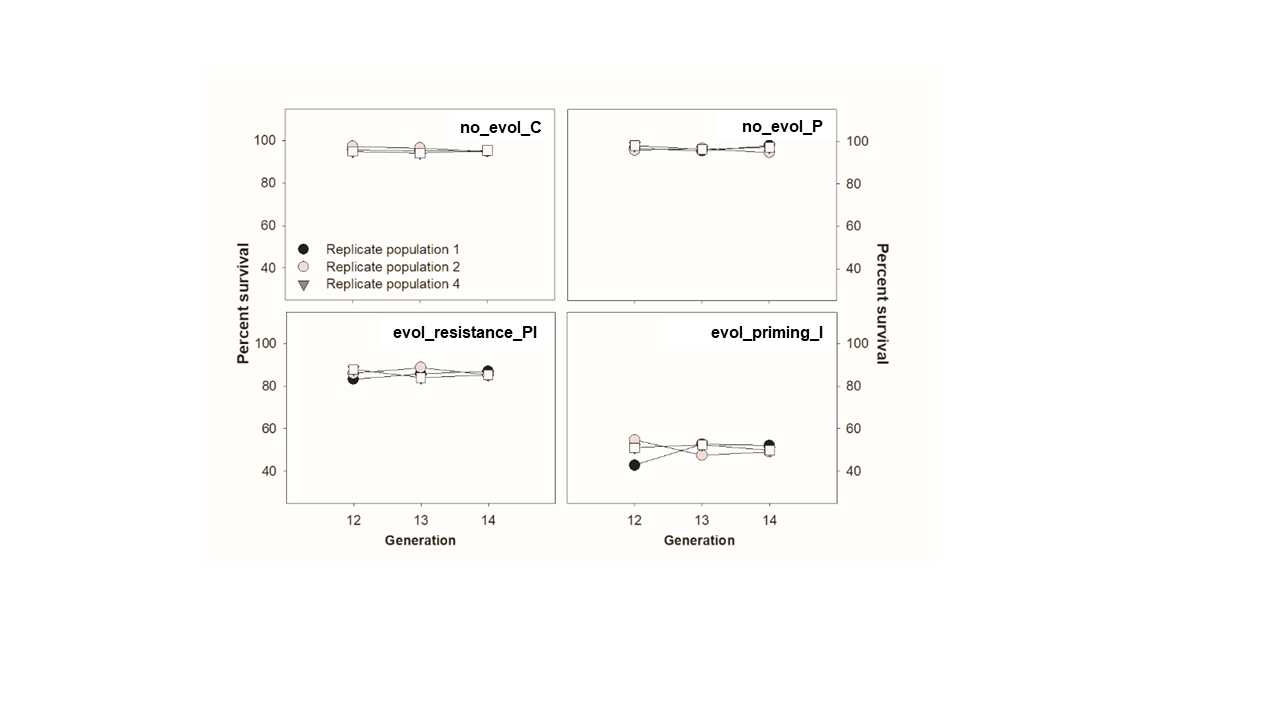

**Figure S4.** Reproductive output of no_evol_C vs. evol_resistance_PI beetles in corn (n=36 females/replicate population/selection regime).

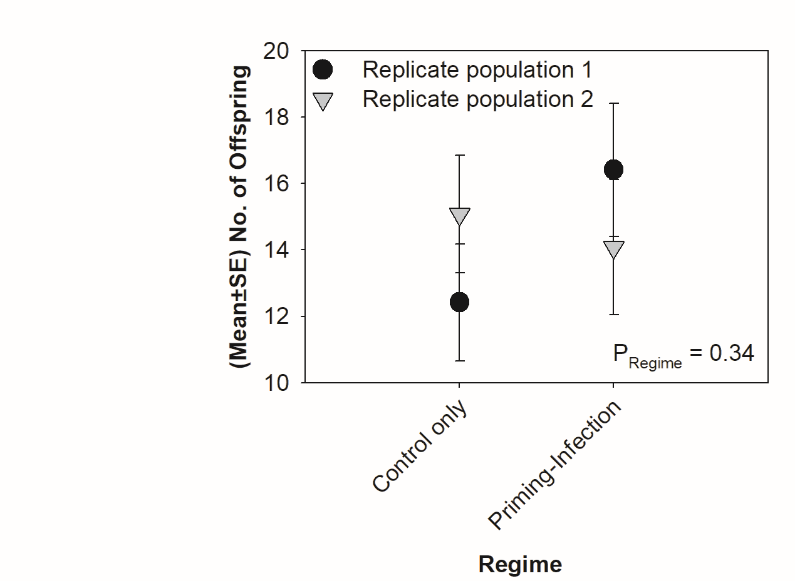

**Figure S5.** Survival curves of naïve males and females under starvation, across selection regimes (*n* = 20 beetles/ sex/ replicate population/ selection regime).

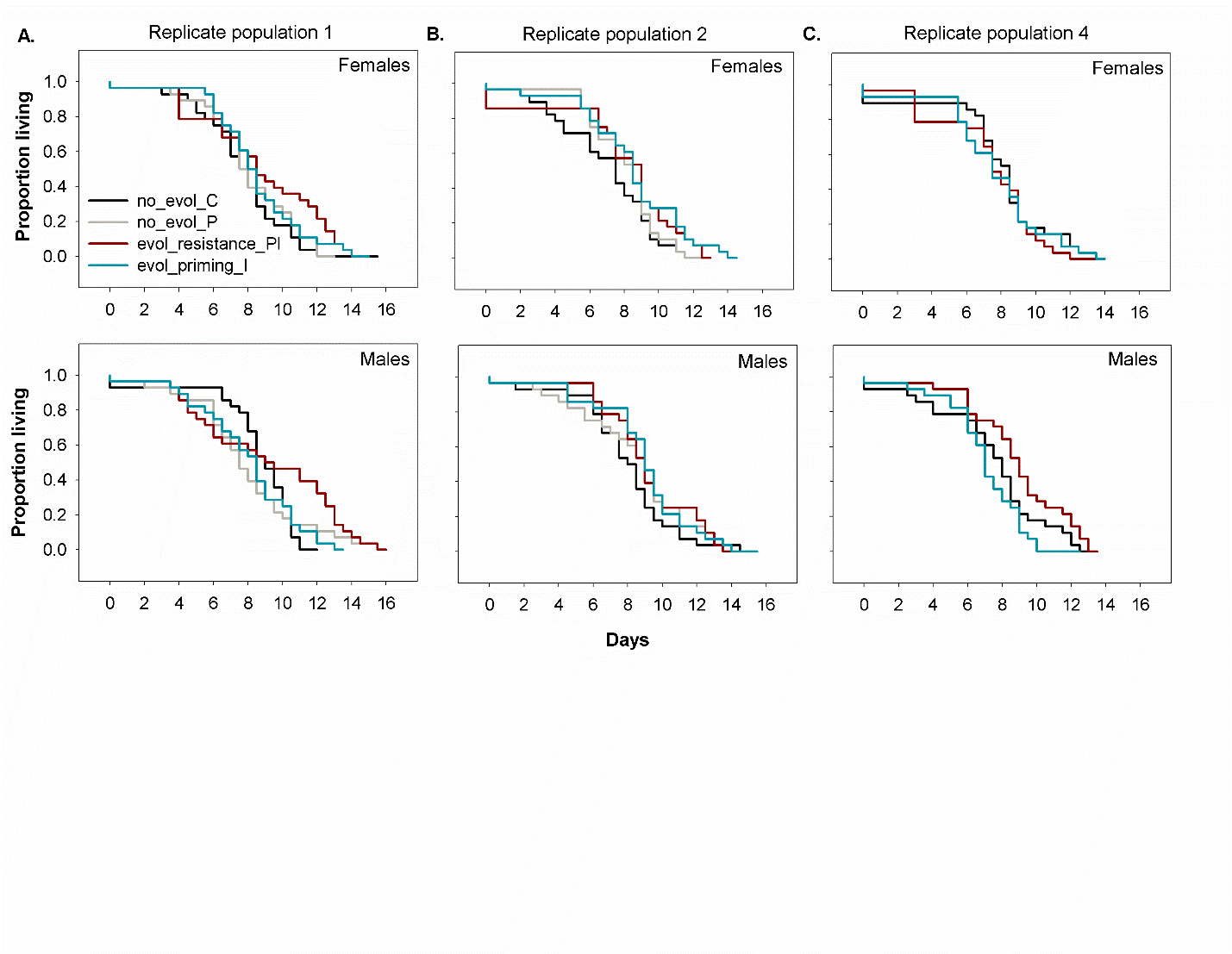

**Figure S6.** Survival curves of naïve females under normal condition up to 95 days post-emergence, across selection regimes (*n* = 48 females/ replicate population/ selection regime).

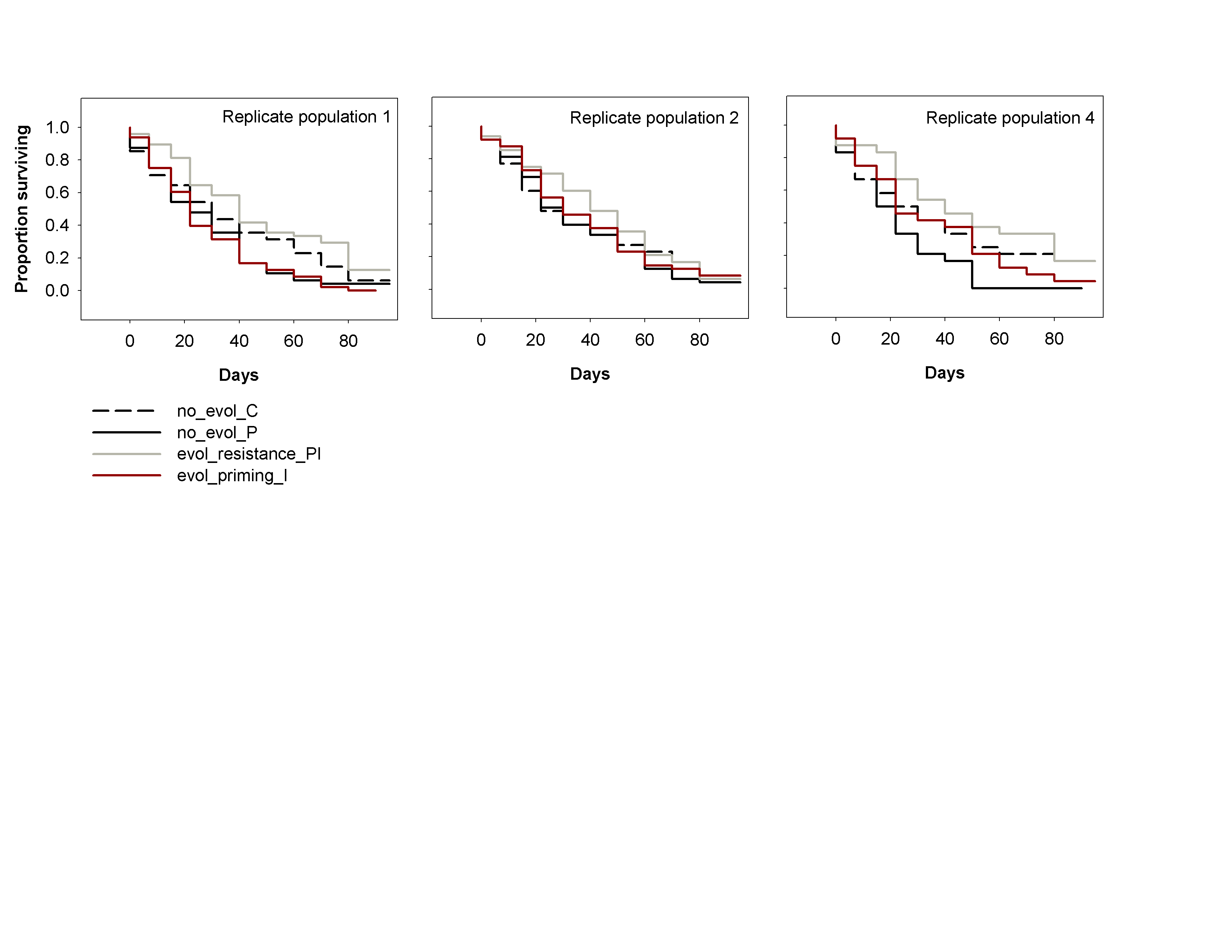

**Figure S7. (A)** Survival curves of the trans-generationally primed evol_priming_I offspring. P values for the impact of TGIP are reported in each panel. **(B)** Survival benefits of WGIP vs. TGIP, based on the hazard function calculated from the survival data of parents (Fig S1 & 2) and offspring. A greater hazard ratio indicates a higher benefit of priming. P values for the impact of priming type (P _Priming_) and sex (P _Sex_) are reported in the panel.

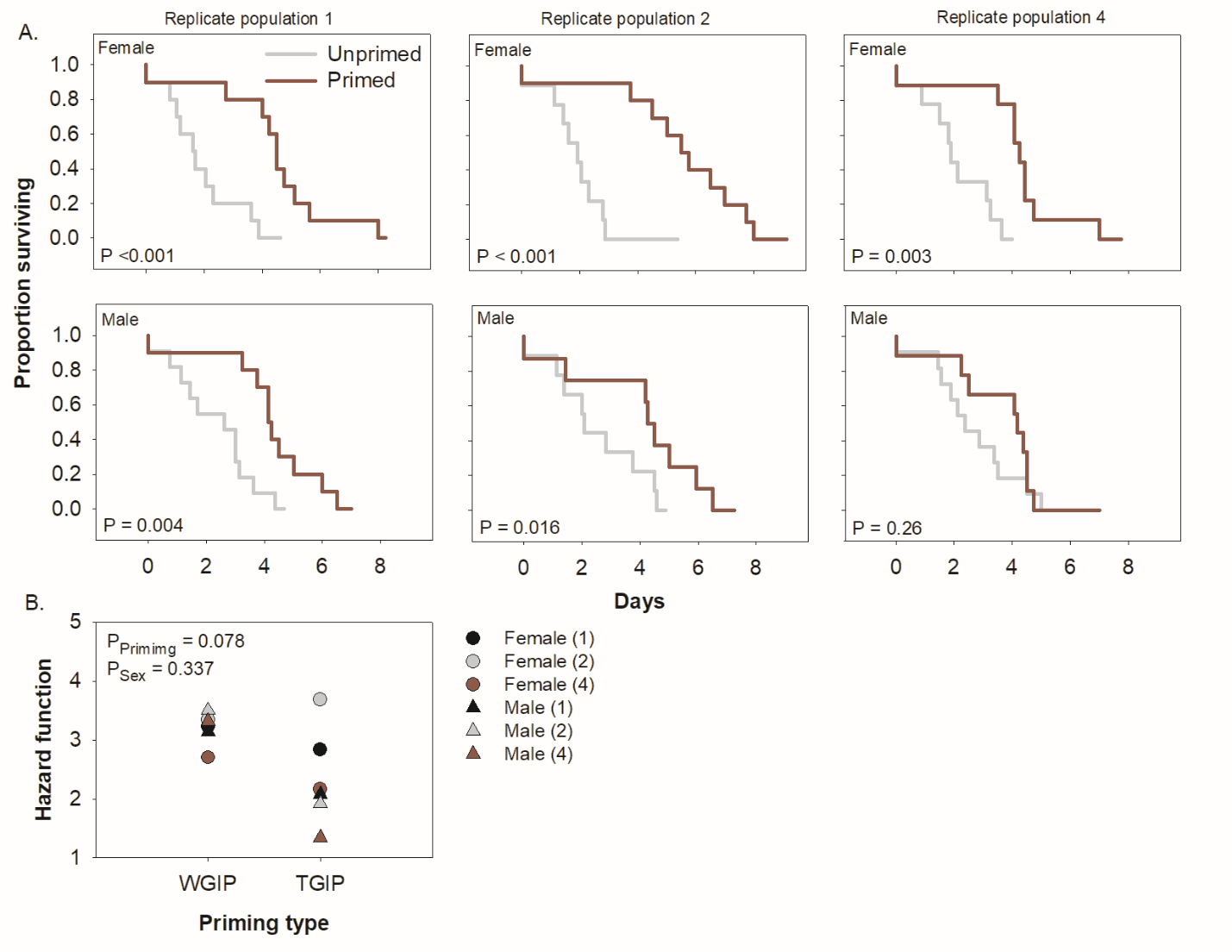

**SUPPLEMENTARY TABLES**

**Table S1**. **(A.)** Number of females that were alive to reproduce after within-generation priming and infection across replicate populations, selection regimes and treatments. **(B.)** Summary of Wilcoxon Rank Sum Test on post-infection replicate size across priming treatments and selection regimes.

| A. | **Replicate population** | **Selection regime** | **Naïve** | **Unprimed** | **Primed** |
| --- | --- | --- | --- | --- | --- |
|  | 1 | no_evol_C | 5 | 7 | 5 |
|  |  | evol_priming_I | 8 | 9 | 10 |
|  |  | no_evol_P | 7 | 11 | 12 |
|  |  | evol_resistance_PI | 6 | 5 | 11 |
|  | 2 | no_evol_C | 7 | 10 | 10 |
|  |  | evol_priming_I | 6 | 8 | 10 |
|  |  | no_evol_P | 8 | 10 | 8 |
|  |  | evol_resistance_PI | 10 | 8 | 10 |
|  | 4 | no_evol_C | 6 | 8 | 9 |
|  |  | evol_priming_I | 6 | 9 | 7 |
|  |  | no_evol_P | 11 | 10 | 10 |
|  |  | evol_resistance_PI | 7 | 8 | 10 |

| **B.** | **Trait** | **χ2** | **df** | **P** |
| --- | --- | --- | --- | --- |
|  | Treatment | 1.545 | 1 | 0.213 |
|  | Regime | 5.452 | 3 | 0.141 |

**Table S2.** **(A)** Summary of mixed effects Cox model, fitting the model to estimate evolved priming response in females. We used data from the naïve infected, unprimed infected and the uninfected control treatments and specified the model as: survival ~ selection regime + treatment + (1|selection regime/ replicate population), with selection regime and treatment as fixed effects, and replicate populations nested within a selection regime as a random effect **(B)** Summary of Cox proportional hazard analysis for each replicate population on survival data for within-generation priming in females.

| **A.** | **Source** | **loglik** | **χ2** | **Df** | **P** |
| --- | --- | --- | --- | --- | --- |
|  | Selection regime | -1887.6 | 14.29 | 3 | **0.002** |
|  | Treatment | -1894.7 | 111.06 | 2 | **>0.001** |
|  | Selection regime × Treatment | -1878.7 | 17.66 | 6 | **0.007** |
|  | *Random effects* | *Std Dev* |  |  |  |
|  | Replicate population | 0.001 |  |  |  |
| **B.** | **Regime** | **Replicate population** | **df** | **χ2** | **P** |
|  | No_evol_C | 1 | 1 | 0.414 | 0.519 |
|  | Evol_priming_I |  | 1 | 6.502 | **0.01** |
|  | No_evol_P |  | 1 | 0.247 | 0.618 |
|  | Evol_resistance_PI |  | 1 | 0.064 | 0.799 |
|  | No_evol_C | 2 | 1 | 0.297 | 0.585 |
|  | Evol_priming_I |  | 1 | 6.493 | **0.01** |
|  | No_evol_P |  | 1 | 1.961 | 0.161 |
|  | Evol_resistance_PI |  | 1 | 0.005 | 0.942 |
|  | No_evol_C | 4 | 1 | 0.088 | 0.765 |
|  | Evol_priming_I |  | 1 | 4.783 | **0.028** |
|  | No_evol_P |  | 1 | 0.721 | 0.395 |
|  | Evol_resistance_PI |  | 1 | 0.032 | 0.855 |

**Table S3.** **(A.)** Summary of mixed effects Cox model, fitting the model to estimate evolved priming response in males. The model has been specified in Table S2. **(B.)** Summary of Cox proportional hazard analysis for each replicate population on survival data for within-generation priming in males.

| **A.** | **Source** | **loglik** | **χ2** | **Df** | | **P** |
| --- | --- | --- | --- | --- | --- | --- |
|  | Selection regime | -1830.9 | 14.095 | | 3 | **0.002** |
|  | Treatment | -1837.9 | 101.81 | | 2 | **<0.001** |
|  | Selection regime × Treatment | -1818.2 | 25.249 | | 6 | **<0.001** |
|  | *Random effects* | *Std Dev* |  | |  |  |
|  | Replicate population | 0.001 |  | |  |  |
| **B.** | **Regime** | **Replicate population** | **df** | | **χ2** | **P** |
|  | No_evol_C | 1 | 1 | 0.743 | | 0.388 |
|  | Evol_priming_I |  | 1 | 7.562 | | **0.005** |
|  | No_evol_P |  | 1 | 1.388 | | 0.238 |
|  | Evol_resistance_PI |  | 1 | 0.759 | | 0.383 |
|  | No_evol_C | 2 | 1 | 0.449 | | 0.502 |
|  | Evol_priming_I |  | 1 | 5.766 | | **0.016** |
|  | No_evol_P |  | 1 | 0.882 | | 0.347 |
|  | Evol_resistance_PI |  | 1 | 0.0002 | | 0.988 |
|  | No_evol_C | 4 | 1 | 0.019 | | 0.888 |
|  | Evol_priming_I |  | 1 | 4.843 | | **0.027** |
|  | No_evol_P |  | 1 | 1.531 | | 0.215 |
|  | Evol_resistance_PI |  | 1 | 0.117 | | 0.731 |

**Table S4.** Summary of Cox proportional hazard analysis on survival data of naïve beetles infected with Bt.

|  | **Comparison** | **Replicate population** | **Hazard function** | **df** | **χ2** | **P** |
| --- | --- | --- | --- | --- | --- | --- |
| Mothers | (no_evol_C vs | 1 | 1.51 | 1 | 0.973 | 0.323 |
|  | no_evol_P) | 2 | 2.34 | 1 | 3.478 | 0.062 |
|  |  | 4 | 1.36 | 1 | 0.519 | 0.471 |
|  | (no_evol_C vs | 1 | **5.87** | 1 | 11.265 | **0.001** |
|  | evol_resistance_PI) | 2 | **4.3** | 1 | 7.882 | **0.005** |
|  |  | 4 | **3.03** | 1 | 5.318 | **0.021** |
|  | (no_evol_C vs | 1 | 0.95 | 1 | 0.01 | 0.919 |
|  | evol_priming_I) | 2 | 1.34 | 1 | 0.504 | 0.473 |
|  |  | 4 | 0.958 | 1 | 0.011 | 0.912 |
| Fathers | (no_evol_C vs | 1 | 3.03 | 1 | 5.391 | **0.02** |
|  | no_evol_P) | 2 | 1.41 | 1 | 0.657 | 0.417 |
|  |  | 4 | 1.28 | 1 | 0.358 | 0.549 |
|  | (no_evol_C vs | 1 | **28.9** | 1 | 20.407 | **<0.001** |
|  | evol_resistance_PI) | 2 | **8.52** | 1 | 13.608 | **0.001** |
|  |  | 4 | **4.37** | 1 | 8.24 | **0.004** |
|  | (no_evol_C vs | 1 | 1.54 | 1 | 1.052 | 0.305 |
|  | evol_priming_I) | 2 | 0.91 | 1 | 0.043 | 0.835 |
|  |  | 4 | 1.15 | 1 | 0.107 | 0.743 |

**Table S5. S**ummary of Wilcoxon Rank Sum test on beetle reproduction before infection as a function of within-generation priming treatment and selection regime.

| **Replicate population** | **Trait** | **χ2** | **df** | **P** |
| --- | --- | --- | --- | --- |
| 1 | Selection Regime | 6.046 | 3 | 0.109 |
|  | Priming Treatment | 1.055 | 2 | 0.589 |
| 2 | Selection Regime | 2.933 | 3 | 0.402 |
|  | Priming Treatment | 1.531 | 2 | 0.464 |
| 4 | Selection Regime | 7.653 | 3 | 0.0537 |
|  | Priming Treatment | 1.647 | 2 | 0.438 |

**Table S6.** Summary of Wilcoxon Rank Sum test for changes in reproductive output after infection across replicate populations and selection regimes.

| **Regime** | **Replicate populations** | **Treatment** | **χ2** | **df** | **P** |
| --- | --- | --- | --- | --- | --- |
| No_evol_C | 1 | Naive | 6.043 | 1 | **0.013** |
|  |  | Unprimed | 8.337 | 1 | **0.003** |
|  |  | Primed | 0.134 | 1 | 0.713 |
|  | 2 | Naive | 6.685 | 1 | **0.009** |
|  |  | Unprimed | 9.486 | 1 | **0.002** |
|  |  | Primed | 5.353 | 1 | **0.02** |
|  | 4 | Naive | 4.854 | 1 | **0.027** |
|  |  | Unprimed | 10.54 | 1 | **0.001** |
|  |  | Primed | 4.71 | 1 | **0.029** |
| No_evol_P | 1 | Naive | 5.55 | 1 | **0.018** |
|  |  | Unprimed | 6.247 | 1 | **0.012** |
|  |  | Primed | 5.225 | 1 | **0.022** |
|  | 2 | Naive | 5.089 | 1 | **0.024** |
|  |  | Unprimed | 1.258 | 1 | 0.262 |
|  |  | Primed | 4.5 | 1 | **0.033** |
|  | 4 | Naive | 0.718 | 1 | 0.396 |
|  |  | Unprimed | 7.321 | 1 | **0.006** |
|  |  | Primed | 0.652 | 1 | 0.419 |
| Evol_resistance_PI | 1 | Naive | 1.121 | 1 | 0.289 |
|  |  | Unprimed | 4.84 | 1 | 0.08 |
|  |  | Primed | 0.067 | 1 | 0.794 |
|  | 2 | Naive | 0.542 | 1 | 0.461 |
|  |  | Unprimed | 0.184 | 1 | 0.667 |
|  |  | Primed | 4.73 | 1 | **0.029** |
|  | 4 | Naive | 0.095 | 1 | 0.757 |
|  |  | Unprimed | 4.898 | 1 | **0.026** |
|  |  | Primed | 1.933 | 1 | 0.164 |
| Evol_priming_I | 1 | Naive | 4.486 | 1 | **0.034** |
|  |  | Unprimed | 8.475 | 1 | **0.003** |
|  |  | Primed | 11.91 | 1 | **0.001** |
|  | 2 | Naive | 11.81 | 1 | **0.001** |
|  |  | Unprimed | 8.408 | 1 | **0.003** |
|  |  | Primed | 6.324 | 1 | **0.011** |
|  | 4 | Naive | 4.484 | 1 | **0.034** |
|  |  | Unprimed | 3.163 | 1 | 0.075 |
|  |  | Primed | 5.66 | 1 | **0.017** |

**Table S7.** Summary of linear mixed effects model on the reproductive output of primed and infected beetles across I. all selection regimes; II. no_evol_C & no_evol_P; III. no_evol_C & evol_resistance_PI; IV. no_evol_C & evol_priming_I. We specified the model as: No. of offspring ~ Priming treatment × Selection regime + (1 ǀ Replicate population)

|  | **Assay** | **Trait** | **df** | **χ2** | **P** |
| --- | --- | --- | --- | --- | --- |
| I. | All selection regimes | Priming treatment (P) | 2 | 6.193 | **0.045** |
|  |  | Selection regime (S) | 3 | 42.11 | **<0.001** |
|  |  | P × S | 6 | 14.687 | **0.022** |
|  |  | *Random effects* | *Std dev* |  |  |
|  |  | Replicate population | 0.000 |  |  |
| II. |  | Priming treatment (P) | 2 | 10.383 | **0.005** |
|  | no_evol_C vs. | Selection regime (S) | 1 | 8.291 | **0.003** |
|  | no_evol_P | P × S | 2 | 2.789 | 0.054 |
|  |  | *Random effects* | *Std dev* |  |  |
|  |  | Replicate population | 0.000 |  |  |
| III. |  | Priming treatment (P) | 2 | 9.278 | **0.009** |
|  | no_evol_C vs. | Selection regime (S) | 1 | 15.206 | **<0.001** |
|  | evol_resistance_PI | P × S | 2 | 11.365 | **0.003** |
|  |  | *Random effects* | *Std dev* |  |  |
|  |  | Replicate population | 0.001 |  |  |
| IV. |  | Priming treatment (P) | 2 | 10.878 | **0.004** |
|  | no_evol_C vs. | Selection regime (S) | 1 | 2.197 | 0.138 |
|  | evol_priming_I | P × S | 2 | 8.058 | **0.017** |
|  |  | *Random effects* | *Std dev* |  |  |
|  |  | Replicate population | 0.000 |  |  |

**Table S8.** Summary of a linear mixed effects model on the reproductive output of naïve (without priming and infection) no_evol_C & evol_resistance_PI beetles reared in corn [Model: No. of offspring~ Selection regime + (1 ǀ Replicate population)]

| **Assay** | **Trait** | **df** | **denDF** | **F-value** | **P** |
| --- | --- | --- | --- | --- | --- |
| Corn | Selection regime | 1 | 2 | 1.595 | 0.34 |
| (no_evol_C vs. | *Random effects* | *Std dev* |  |  |  |
| evol_resistance_PI) | Replicate population | 0.003 |  |  |  |

**Table S9**. Summary of mixed effects Cox model analysis on survival under starvation (of unhandled naïve males and females), with reduced output. Survival ~ Selection regime + Sex + (1| Selection regime/ Replicate population)

| **Trait** | **loglik** | **df** | **χ2** | **P** |
| --- | --- | --- | --- | --- |
| Selection regime | -3339.8 | 3 | 10.156 | **0.01** |
| Sex | -3337.9 | 1 | 3.752 | 0.052 |
| *Random effects* | *Std Dev* |  |  |  |
| Replicate population | 0.075 |  |  |  |

**Table S10**. Summary of mixed effects Cox model analysis on long-term survival of unhandled naïve females [Model: Survival ~ Selection regime + (1|Selection regime/ Replicate population)]

| **Trait** | **loglik** | **df** | **χ2** | **P** |
| --- | --- | --- | --- | --- |
| Selection regime | -2399.2 | 3 | 20.313 | **<0.001** |
| *Random effects* | *Std Dev* |  |  |  |
| Replicate population | 0.002 |  |  |  |

**Table S11.** Summary of linear mixed effects model on early survival and developmental rate [Model: Trait ~ Selection regime + (1|Replicate population)]

|  | **Experiment** | **Trait** | **χ2** | **Df** | **P-value** |
| --- | --- | --- | --- | --- | --- |
| A. | Egg-hatchability | Regime | 20.494 | 2 | **<0.001** |
|  | *Random effects* | *Std Dev* |  |  |  |
|  | Replicate population | 0.000 |  |  |  |
| B. | Fraction of pupae | Regime | 5.084 | 2 | 0.08 |
|  | *Random effects* | *Std Dev* |  |  |  |
|  | Replicate population | 0.02 |  |  |  |
| C. | Fraction of adults | Regime | 9.639 | 2 | **0.008** |
|  | *Random effects* | *Std Dev* |  |  |  |
|  | Replicate population | 0.000 |  |  |  |
| D. | Percent viability | Regime | 28.597 | 2 | **<0.001** |
|  | *Random effects* | *Std Dev* |  |  |  |
|  | Replicate population | 0.000 |  |  |  |

**Table S12.** Summary of the linear mixed effects model on mean post-infection survival of offspring as a function of selection regime and parental priming status (with reduced output). [Model: Mean post-infection survival~ Selection regime × Parental priming + (1ǀReplicate population)]

| **Trait** | **χ2** | **Df** | **P-value** |
| --- | --- | --- | --- |
| Selection regime | 601.034 | 3 | **<0.001** |
| Parental priming | 35.211 | 2 | **<0.001** |
| Selection regime × Parental priming | 74.02 | 6 | **<0.001** |
| *Random effects* | *Std Dev* |  |  |
| Replicate population | 0.05 |  |  |

**Table S13.** Summary of Cox proportional hazard analysis on offspring survival from evol_priming_I regime as a function of parental priming status. [Model: Survival ~ Parental priming status + (1| parental mating pair/ replicate offspring)]

| Replicate population | Sex | Hazard function | df | **χ2** | P |
| --- | --- | --- | --- | --- | --- |
| 1 | Female | **2.83** | 1 | 13.764 | **<0.001** |
|  | Male | **2.078** | 1 | 8.412 | **0.004** |
| 2 | Female | **3.67** | 1 | 19.764 | **<0.001** |
|  | Male | **1.919** | 1 | 5.68 | **0.016** |
| 4 | Female | **2.16** | 1 | 9.057 | **0.003** |
|  | Male | 1.3 | 1 | 1.25 | 0.26 |

**Table S14.** Summary of Wilcoxon Rank Sum test on progeny fitness before infection, as a function of selection regime and priming treatment.

| **Block** | **Trait** | **χ2** | **df** | **P** |
| --- | --- | --- | --- | --- |
| 1 | Regime | 7.792 | 3 | 0.052 |
|  | Treatment | 2.147 | 1 | 0.143 |
| 2 | Regime | 3.499 | 3 | 0.32 |
|  | Treatment | 0.418 | 1 | 0.517 |
| 4 | Regime | 6.867 | 3 | 0.077 |
|  | Treatment | 0.029 | 1 | 0.864 |

**Table S15:** Summary of Wilcoxon Rank Sum test for changes in offspring’s reproductive output after infection across replicate populations and selection regimes.

| **Replicate population** | **Treatment** | **Selection regime** | **χ2** | **df** | **P** |
| --- | --- | --- | --- | --- | --- |
| 1 | Unprimed | no_evol_C | 3.303 | 1 | **0.0691** |
|  | Primed |  | 7.828 | 1 | **0.005** |
|  | Unprimed | no_evol_P | 0.157 | 1 | 0.691102 |
|  | Primed |  | 4.3246 | 1 | **0.037563** |
|  | Unprimed | evol_resistance_PI | 1.107 | 1 | 0.292 |
|  | Primed |  | 0.918 | 1 | 0.337 |
|  | Unprimed | evol_priming_I | 10.115 | 1 | **0.001** |
|  | Primed |  | 0.894 | 1 | 0.344 |
| 2 | Unprimed | no_evol_C | 3.481 | 1 | **0.062** |
|  | Primed |  | 3.951 | 1 | **0.046** |
|  | Unprimed | no_evol_P | 0.903 | 1 | 0.341 |
|  | Primed |  | 4.698 | 1 | **0.03** |
|  | Unprimed | evol_resistance_PI | 0.707 | 1 | 0.4 |
|  | Primed |  | 1.949 | 1 | 0.162 |
|  | Unprimed | evol_priming_I | 5.080 | 1 | **0.024** |
|  | Primed |  | 9.163 | 1 | **0.002** |
| 4 | Unprimed | no_evol_C | 5.08 | 1 | **0.024** |
|  | Primed |  | 6.33 | 1 | **0.011** |
|  | Unprimed | no_evol_P | 2.668 | 1 | 0.102 |
|  | Primed |  | 5.512 | 1 | **0.018** |
|  | Unprimed | evol_resistance_PI | 1.106 | 1 | 0.292 |
|  | Primed |  | 0.148 | 1 | 0.7 |
|  | Unprimed | evol_priming_I | 9.29 | 1 | **0.002** |
|  | Primed |  | 8.235 | 1 | **0.004** |

**Table S16.** Summary of best reduced linear mixed effects model on offspring fitness after infection. We specified the model as: Mean no. of offspring ~ Parental priming status + Selection regime + (1 ǀ Replicate population)

| **Traits** | **χ2** | **df** | **P** |
| --- | --- | --- | --- |
| Parental priming status | 4.357 | 1 | **0.036** |
| Selection regime | 28.174 | 3 | **<0.001** |
| *Random effects* | *Std Dev* |  |  |
| Replicate populations | 0.000 |  |  |
